# Comprehensive approach to study branched ubiquitin chains reveals roles for K48-K63 branches in VCP/p97-related processes

**DOI:** 10.1101/2023.01.10.523363

**Authors:** Sven M. Lange, Matthew R. McFarland, Frederic Lamoliatte, Dominika Kwaśna, Linnan Shen, Iona Wallace, Isobel Cole, Lee A. Armstrong, Axel Knebel, Clare Johnson, Virginia De Cesare, Yogesh Kulathu

**Affiliations:** MRC Protein Phosphorylation and Ubiquitylation Unit, University of Dundee, Dundee, Scotland, UK; Department of Biological Chemistry and Molecular Pharmacology, Harvard Medical School, Boston, MA, USA; Malopolska Centre of Biotechnology, Krakow, Poland

**Keywords:** Branched ubiquitin, nanobody, DNA damage response, Deubiquitinase, heterotypic ubiquitin, protein unfolding, protein engineering, protein-protein interaction, ubiquitin binding domain, signal transduction, VCP/p97

## Abstract

Branched ubiquitin (Ub) chains make up a significant proportion of Ub polymers in human cells and are formed when two or more sites on a single Ub molecule are modified with Ub creating bifurcated architectures. Despite their abundance, we have a poor understanding of the cellular functions of branched Ub signals that stems from a lack of facile tools and methods to study them. Here we develop a comprehensive pipeline to define branched Ub function, using K48-K63-branched chains as a case study. We discover branch-specific binders and, by developing a method that monitors cleavage of linkages within complex polyUb, we discover the VCP/p97-associated ATXN3, and MINDY family deubiquitinases to act as debranching enzymes. By engineering and utilizing a branched K48-K63-Ub chain-specific nanobody, we reveal roles for these chains in VCP/p97-related processes. In summary, we provide a blueprint to investigate branched Ub function that can be readily applied to study other branched chain types.

**Highlights:**

- Assembly of defined branched ubiquitin chains enables identification of specific binding proteins
- Development of quantitative DUB assay monitoring cleavage of individual Ub linkages within complex ubiquitin chains identifies debranching enzymes
- Engineering specific, high-affinity nanobody against branched K48-K63 ubiquitin reveals roles in VCP/p97 related processes and DNA damage responses
- General blueprint of new methods and tools for in-depth characterization of branched ubiquitin chains and their underlying biology

## Introduction

The post-translational modification of protein substrates with ubiquitin (Ub) plays a critical role in every major signaling pathway in humans. A cascade of E1, E2 and E3 enzymes catalyzes the conjugation of the C-terminal glycine of Ub to substrate Lys residues or to other Ub molecules via the N-terminal Met or one of seven Lys residues in Ub. Diverse Ub architectures are assembled in this way, ranging from single ubiquitin (monoUb) to complex chains (polyUb) of homo- or heterotypic nature, i.e., containing a single or multiple linkage types in the same Ub chain, respectively (Komander and Rape, 2012). K48- and K63-linkages are the most abundant linkage types found in cells and are well studied individually (Komander and Rape, 2012). Homotypic K48-linked Ub chains have been identified to function mainly in degradative roles by marking substrates for proteasomal degradation. In contrast, homotypic K63-linked polyUb play critical roles during endocytosis, DNA damage repair and innate immune responses.

Sophisticated mass-spectrometric studies have found that a substantial amount of K48- and K63-linkages co-exist in heterotypic, branched chains and these branched K48-K63-Ub chains have been connected to NF-κB signaling and proteasomal degradation (Ohtake et al., 2018, 2016). Such branched architectures are formed when two Ub molecules are attached to Lys residues of a third Ub. Theoretically, 28 different branched Ub architectures can be formed in this way and importantly they account for around 10% of all polyUb formed in human cells (Swatek et al., 2019). Two other branched Ub chain types, K11-K48 and K29-K48, have been detected in cells with roles for them attributed to several processes including protein quality control, ERAD, cell cycle and protein degradation (Haakonsen and Rape, 2019; Ohtake, 2022). In addition, chemically induced degradation of neo-substrates via PROTAC approaches have identified the formation of branched Ub chains (Akizuki et al., 2022). Hence, branched chains may have several important hitherto unappreciated roles. A major limitation to our understanding the function of branched chains is the lack of tools and methods to study them. Huge advances in elucidating the roles of K11-K48-branched chains was made possible by the development of a bispecific antibody that recognizes these chains thereby enabling their detection in cells (Yau et al., 2017). However, such tools for facile detection do not exist for other branched types and requires the use of sophisticated mass spectrometry-based approaches or expression of Ub mutants in cells.

Ubiquitin modifications are recognized by ubiquitin-binding domains (UBDs), which often bind selectively to a specific linkage type. UBDs are structurally diverse and found in a wide range of proteins throughout the ubiquitin system, including signal transducers, ligases and DUBs (Husnjak and Dikic, 2012). While UBDs recognizing polyUb of certain linkage types have been identified (Zhang et al., 2017), specific binders to branched Ub have not been identified to date. One receptor of ubiquitylated substrates is the unfoldase Valosin-containing protein (VCP)/p97 (also known as TERA or Cdc48 in yeast) which is highly conserved in eukaryotes and facilitates unfolding or extraction of its targets. VCP/p97 is a hexameric AAA+-ATPase that is critical in many ubiquitin-dependent pathways. It can function in degradative roles in the proteasomal or lysosomal destruction of proteins or in non-degradative, regulatory roles by extracting proteins from macromolecular complexes or membranes (van den Boom and Meyer, 2017). More than 30 cofactors that directly or indirectly bind to VCP/p97 have been identified, many of which contain UBDs and may function as substrate adapters (Ahlstedt et al., 2022). Some of the major VCP/p97 substrate adapters such as UFD1-NPLOC4, UBXD1, and p47 are thought to function in a mutually exclusive manner, but each to associate with additional cofactors (Buchberger et al., 2015; Meyer and Weihl, 2014). However, mechanistic details on how VCP/p97 and its adapters are implicated in diverse cellular signaling pathways, e.g. how specific ubiquitylated substrates are chosen, are not well understood.

An array of ~100 deubiquitinases (DUBs) cleave Ub linkages thereby fine-tuning or removing Ub signals (Lange et al., 2022). In humans, these are divided into seven families, six being cysteine proteases, and a sole family of metallo-proteases. DUBs play important regulatory roles, and their deregulation has been associated with several diseases including cancer, inflammation and neurodegeneration. Importantly, DUBs can discriminate between polyUb of different linkage types and employ different mechanisms to achieve substrate recognition. For instance, the JAMM, OTU, MINDY and ZUP1 family DUBs cleave polyUb in a linkage selective manner whereas the USP family enzymes are very linkage promiscuous (Abdul Rehman et al., 2021; Faesen et al., 2011; Kwasna et al., 2018; Ritorto et al., 2014). The Josephin family DUBs show different substrate preferences with some of them working as esterases (De Cesare et al., 2021). Of the Josephin family DUBs, ATXN3 and ATXN3L prefer cleaving long polyUb chains albeit with very poor efficiency (Winborn et al., 2008). Indeed, it is recently being increasingly appreciated that Ub chain length in addition to linkage type is a determinant of DUB activity (Hermanns et al., 2018; Kwasna et al., 2018; Mevissen et al., 2013). Moreover, the recent identification of UCHL5 as an enzyme that debranches Ub chains at the proteasome suggests that DUBs may also prefer bifurcated polyUb architectures (Deol et al., 2020; Song et al., 2021).

In this study, we outline a comprehensive strategy to understand the cellular roles of K48-K63-branched Ub chains - identifying binding proteins or ‘reader’ modules, debranching DUBs or ‘eraser/editor’ modules, and developing tools to monitor proteins modified with branched Ub in cells. The basis of our work is the development of a method for enzymatically assembling complex Ub chain architectures in vitro, which enables us to produce well-defined Ub chains containing multiple linkage types. These custom-made Ub chains incorporate immobilization tags or Ub moieties of specific masses, enabling identification of specific binders in cells and the development of a quantitative DUB assay that can identify linkages cleaved within complex chain architectures. Finally, we engineer a nanobody that specifically binds to branched K48-K63-Ub chains with picomolar affinity and use it to reveal roles for these branched chains in VCP/p97-related processes. The methods and pipeline we describe here can be readily applied to reveal the molecular players and functions of other Ub chain architectures.

## Results

### Standardized nomenclature for branched Ub chains

The minimal branched Ub chain unit is typically viewed to be made up of three Ub moieties, with two distal Ub moieties attached to a single proximal Ub and branching potentially creates or disrupts interfaces for protein interactions. However, one can envisage branched tetrameric Ub (Ub4) to be formed when a single Ub branches off a homotypic tri-meric Ub (Ub3) chain as the ‘trunk’ (**Figure 1A**). Importantly, branched Ub4 chains contain additional unique interfaces missing in the minimal branched Ub3 and may therefore encode extra information dependent not only on the type but also the order of linkages from trunk to branch. Hence, we decided to study tetrameric branched Ub since important information may be lost when studying shorter branched Ub3.

**Figure 1.**
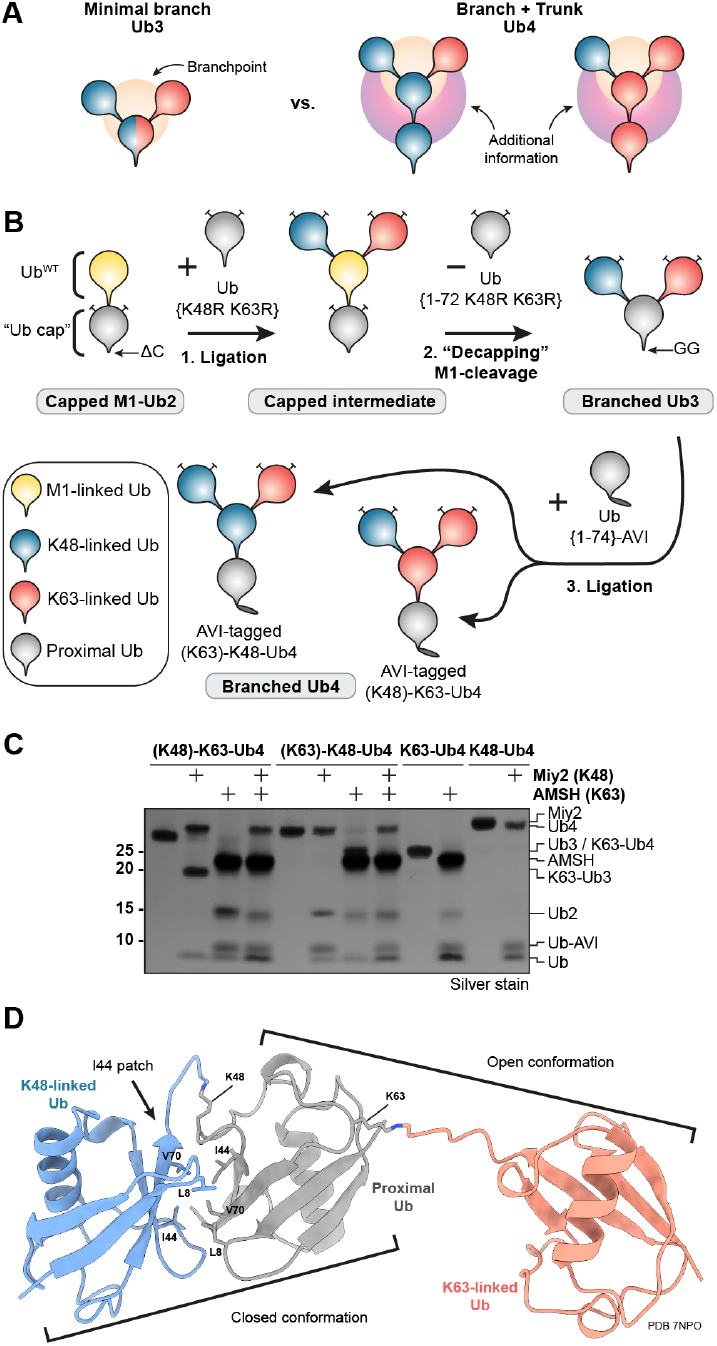
‘Ub capping’ enables ligation of tetrameric branched Ub chains and crystal structure of branched K48-K63-Ub3. A) Schematic depicting differences in binding interfaces of minimal branched triUb (orange) chain in comparison to branched tetraUb (orange and purple). B) Schematic workflow of “Ub-capping” approach for ligation of functionalized, branched K48-K63-Ub4 chains. C) DUB assay with linkage-specific enzymes Miy2/Ypl191c (K48) and AMSH (K63) as quality control for synthesized branched and unbranched Ub4. D) Crystal structure of K48-K63-Ub3 in cartoon representation with K48-linked Ub in blue, K63-linked Ub in red and proximal Ub in grey. Atoms of the I44 interface between K48-linked Ubs and isopeptide linkages are shown as stick models.

The complexity of different chain types and the common use of Ub mutants in the *in vitro* generation of branched chains necessitates a definitive and universal nomenclature to accurately describe the architecture of the Ub chain, as previously proposed terminologies were often limited to short, trimeric branched and mixed chains (Nakasone et al., 2013; Song et al., 2021). We therefore devised a universal nomenclature for chains made of Ub and Ubiquitin-like proteins (UBLs) that is derived from the IUPAC Nomenclature of organic chemistry: The proximal Ub/UBL is the origin of the chain, and the longest chain forms the parent chain (or, in the case of equally long branches, the linkage type carrying the highest number would be denoted the parent chain, e.g. K63 would be parent over K48). The Ub/UBLs of the parent chain are numbered starting at the origin and appending Ub/UBL chains are listed in brackets with a prefix of the substituent’s location on the parental chain from low to high. The total sum of Ub/UBL components is given in square brackets and followed by a list of modifications (e.g. truncations or mutations) in curly brackets for each Ub/UBL moiety (**Figure S1A**). We also envision a simplified alias to be given to each chain in a context-dependent manner to improve readability. For example, the full name of a branched trimer containing K48- and K63-linkages would be 1-(K48-Ub)-K63-Ub2 [Ub3] and the same assembled from Ub^1-72^ and Ub^K48R K63R^ would be 1-(K48-Ub)-K63-Ub2 [Ub3] {1-72, K48R K63R, K48R K63R} and a short alias may be K48-K63-Ub3^1-72^. A mixed Ub chain on the other hand would be denoted for instance as 2-(K48-Ub2)-K63-Ub2. Given the broad familiarity with basic organic chemistry and IUPAC nomenclature among researchers, this nomenclature therefore provides a comprehensive scheme to accurately describe the intrinsic details of Ub chains while also providing intuitive readability. Since investigating heterotypic ubiquitin chains is a rapidly expanding field, we believe that the timely adoption of standardized nomenclature proposed here will avoid future confusion.

### Strategies for the *in vitro* generation of well-defined branched chains

The *in vitro* study of branched and mixed ubiquitin chains containing multiple linkage types is hindered by the inability to generate sufficient quantities of polyUb of defined architectures. Various approaches from chemical synthesis and sortase-based ligation have been devised to generate Ub chains of particular design (El Oualid et al., 2010; Fottner et al., 2019). However, these methods often introduce synthetic linkages in place of the native isopeptide bond, lack scalability, accessibility and the economical nature of enzymatic approaches. To circumvent this, we employed two complementary approaches of enzymatic Ub chain assembly strategies, permitting the assembly of well-defined long and complex Ub chains. Both strategies combine the use of linkage-specific ligation enzymes with Ub mutants that together determine which Ub C-terminal carboxyl group is ligated to which Lysine’s amine. The “Delta-C” approach uses a Ub moiety lacking the C-terminal GlyGly motif (Ub^ΔC^) required for ligation either by C-terminal truncation or replacement of the C-terminus with a functionalized tag (**Figure S1B**) and has previously been applied in similar form to generate branched Ub3 (Deol et al., 2020; Song et al., 2021; Yau et al., 2017). For example, for the assembly of branched K48-K63-Ub3, the UbΔC moiety is mixed with UbK48R K63R and added to a Ub ligation reaction containing K48- and K63-linkage specific E2 enzymes. The Ub3 product is subsequently separated from unreacted substrate and enzymes using cation exchange chromatography. However, a disadvantage of this approach is that the resulting chain lacks the native C-terminus on the proximal Ub which may be important for protein interactions. Moreover, branched tetraUb assembly is not possible as the lack of the native C-terminal residues also prevents further chain extension.

To circumvent this problem, we developed the “Ub-capping” strategy which permits the assembly of longer and more complex chains (**Figure 1B, S1C**). The core idea of this technique is the use of a “capped” M1-linked diUb (M1-Ub2) harboring Lys-to-Arg mutations and a truncated C-terminus on the proximal Ub (the “cap”). Only Lys residues of the distal Ub of this capped Ub2 are therefore available for ligation with other Ub C-termini. In the case of branched K48-K63-Ub chains, we use Ub^K48R K63R^ and site-specific E2 enzymes to attach distal K48- and K63-linked Ubs to the capped M1-Ub2. Using the nomenclature described above, the intermediate chain formed during the Ub-capping approach shown in Fig 1 with a full name of 2-(K48-Ub)-2-(K63-Ub)-M1-Ub2 [Ub4] {1-72 K48R K63R, wt, K48R K63R, K48R K63R} or short alias of (K48)-(K63)-M1-Ub4^CAP^ (**Figure S1A**). Subsequently, the proximal cap can be removed using the M1-specific DUB OTULIN. Importantly, this “decapping” step exposes a native C-terminus on the now proximal Ub that is then available for further ligation steps. Using this approach, we have successfully assembled milligram quantities of pure K48-K63-branched Ub chains and the linkage composition of the assembled chains was confirmed using the K48- and K63-linkage specific DUBs Ypl191c/Miy2 and AMSH (**Figure 1C**).

To further establish the purity of the assembled branched chains and to get insights into the structure of K48-K63 branched chains, we crystallized K48-K63-Ub3. The structure was determined using molecular replacement and refined to the statistics shown in **Table S1**. In the structure of K48-K63-Ub3, the K48-linked Ubs adopt a closed conformation with interactions between the I44 patches of the two Ub moieties (**Figure 1D**), while the K63-linked Ubs adopt an open, extended conformation. These closed and open configurations have been observed previously for K48- and K63-linked Ub2, respectively (Cook et al., 1992; Weeks et al., 2009), and for branched and mixed K48-K63-Ub3 in NMR analyses (Nakasone et al., 2013).

### Identification of linkage-specific binders of homotypic and branched Ub chains

How a posttranslational modification is decoded can reveal how it is utilized in transducing a signal (Pawson, 2004). To identify cellular proteins that bind to branched K48-K63-Ub chains, we utilized the Ub-capping approach to generate branched and unbranched K48- and K63-linked tetraUb (Ub4) that incorporated a SpyTag (Keeble et al., 2017) on the C-terminus of the proximal Ub for covalent and site-directed immobilization of chains on SpyCatcher-coupled agarose beads (**Figure 2A, S2A**). Importantly, immobilization via a defined anchor ensures that the branched interfaces are available for protein interaction. Next, the immobilized chains were incubated in quadruplicate with protease inhibitor-treated U2OS cell lysates, and bound proteins subjected to data-independent acquisition mass spectrometry (DIA MS/MS) (**Figure 2A, S2B**). We analyzed the normalized binding Z-scores of a total of 7999 unique protein isoforms identified across the chain pulldown samples and found 130 proteins with Z-scores differing significantly to at least one other chain type (**Figure 2B, Supplementary Data**).

**Figure 2.**
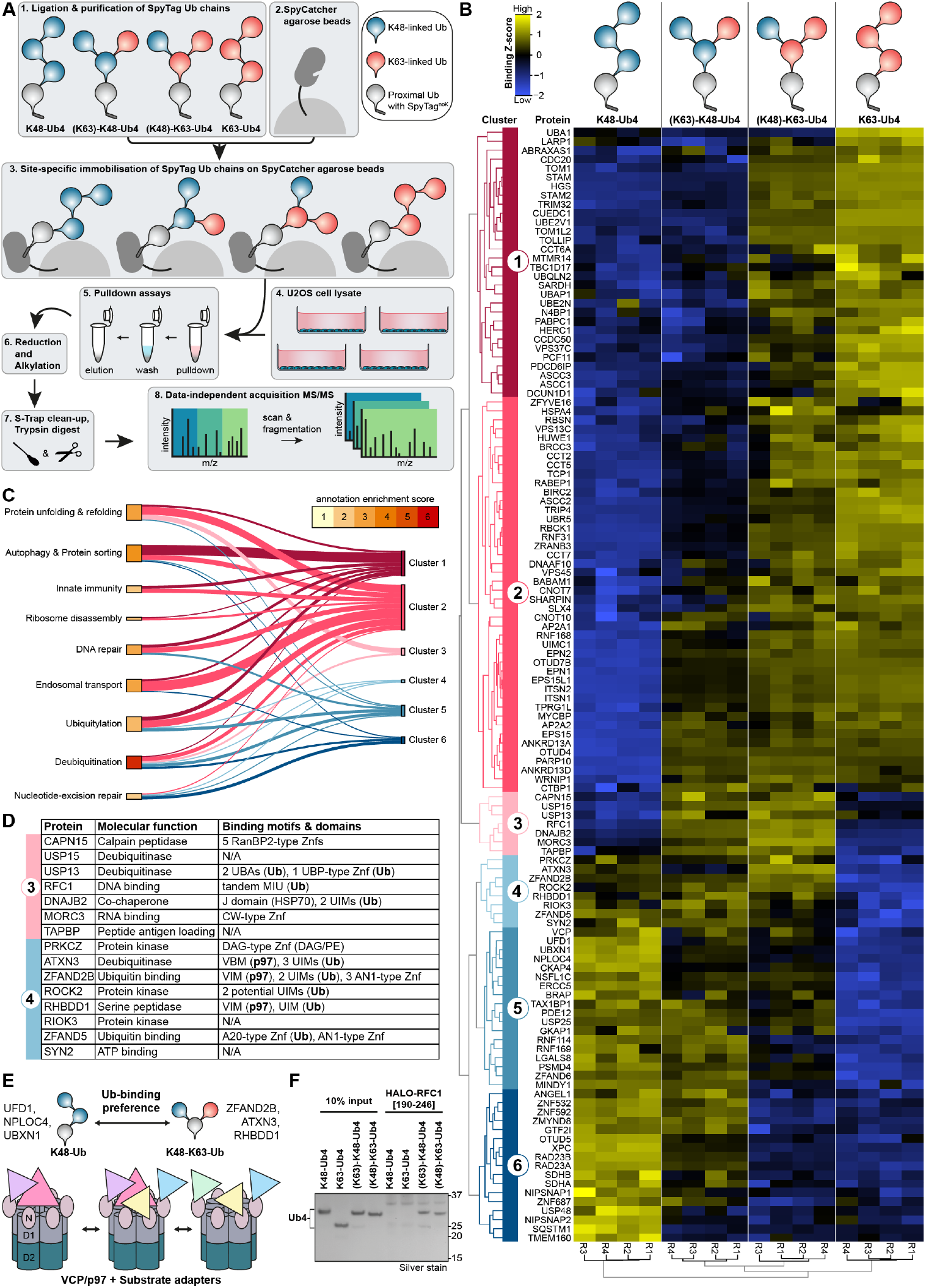
Identification of branched K48-K63-Ub chain-specific binders. (Continued on next page) A) Schematic workflow of pulldown from U2OS cell lysates using functionalized Ub4 chains and subsequent MS-MS & DIA analysis. B) Heatmap showing binding Z-scores of 130 proteins with statistically significant differences in binding profiles identified in quadruplicate Ub chain pulldowns. Schematics of chains used in pulldown are depicted on top. Spatial Euclidean distance computations were applied to rank proteins (left tree) and replicates (bottom tree). The six main distance clusters of proteins representing binding preferences are color-coded from red to blue. C) Sankey diagram connecting the six distance clusters to annotation clusters of DAVID gene ontology analysis colored by annotation enrichment score. D) Table of molecular functions and known binding motifs of proteins specifically associated with branched K48-K63-Ub chains (Clusters 3 and 4). E) Schematic of VCP/p97 sub-complexes with varying Ub chain binding preferences. F) Silver-stained SDS-PAGE analysis of HALO pulldown with recombinant HALO-tagged RFC1 tandem MIU [190-246] and branched/unbranched Ub4 containing K48- and K63-linkages.

Sorting the 130 significant hits by spatial Euclidean distance computations classified the proteins into six main clusters of Ub chain interactors: proteins that mainly bind unbranched K63 chains (Clusters 1 and 2), branched K48-K63 chains (Clusters 3 and 4) and unbranched K48 chains (Clusters 5 and 6) (**Figure 2B**). Gene ontology enrichment analysis reveals a strong association with biological processes that have previously been shown to be ubiquitin-dependent, with the strongest annotation enrichment found for deubiquitination, which was present across all six clusters (**Figure 2C**).

Cluster 1 contains 30 proteins, which associate preferentially with long K63-linked Ub chains (>2 Ub), but not with K48-linked Ub or the K63-linked Ub that is branched-off a K48-linked trunk. In contrast, the 44 proteins in Cluster 2 show increasing propensity to interact with the shorter K63-diUb present in this branched chain. Proteins in these two K63-linkage binding clusters are strongly linked to biological processes previously connected to K63-linked ubiquitylation, including protein unfolding and refolding, autophagy and protein sorting, and endosomal transport (**Figure 2C**). These include previously annotated K63-binding proteins such as the BRCA1-complex subunit ABRAXAS-1, K63-specific DUB BRCC36, ZRANB3, UIMC1/RAP80 (Cooper et al., 2009; Patterson-Fortin et al., 2010; Sims and Cohen, 2009), trafficking-related proteins EPS15, ANKRD13D, TOM1, STAM, STAM2, TOM1, HGS, TOM1L2 and TOLLIP (Erpapazoglou et al., 2012; Lauwers et al., 2009; Messick and Greenberg, 2009; Nathan et al., 2013; Sobhian et al., 2007). In addition, we also identified less studied proteins with annotated UBDs like CUEDC1 and ASCC2 (CUE), N4BP1 (CoCUN) (Gitlin et al., 2020; Nepravishta et al., 2019), CCDC50 (MIU) (Hou et al., 2021) and RBSN (UIM).

A further 18 proteins in Cluster 5 mainly bind to K48-linkages in chains irrespective of whether they are within homotypic or branched architectures. Similarly, Cluster 6 contains 17 proteins that show strong binding to unbranched K48-chains and weak association with 2-(K63)-K48-Ub4, suggesting they prefer longer K48-chains (>2 Ub) or cannot bind to single K48-linked Ub branching off a K63-linked trunk. Proteins primarily binding to the unbranched K48-linked chains include the proteasomal ubiquitin-binding component PSDM4/Rpn10, the segregase VCP/p97 and its substrate adapters UBXN1, UFD1, NSFL1C and NPLOC4 (**Figure 2B**). Other known K48 binders present in this cluster include the proteasome shuttling factors RAD23A and RAD23B, the DUBs MINDY1, OTUD5, USP25 and USP48 (Abdul Rehman et al., 2021; Beck et al., 2021; Raasi et al., 2004). Of note, other proteasomal subunits were identified across all chain pulldowns, but not significantly enriched by a specific chain, indicating that these subunits can associate with long polyUb chains independent of linkage type (**Supplementary Data**). We also identify several proteins with no annotated UBDs such as MTMR14, ZFAND6 and TBC1D17 as binders to K48- and K63-linked chains. However, it is unclear if these proteins bind directly to the Ub chains or if they were identified as part of larger macromolecular complexes which contains a UBD.

Strikingly, we identified 7 proteins (Cluster 3) that strongly associate with the two branched chain architectures used in the pulldown, but not with the unbranched K48- or K63-linked chains (**Figure 2B**). This cluster contains proteins that have previously been implicated in DNA replication (RFC1), Histone deubiquitination (USP15), Histone methylation reading (MORC3), ERAD (USP13, DNAJB2), and peptide antigen loading (TAPBP) (**Figure 2D**) (Chapple and Cheetham, 2003; Gaidt et al., 2021; Li et al., 2000; Long et al., 2014; Podust et al., 1995; Zhang et al., 2019). A further 8 proteins in Cluster 4 display high Z-scores for the two branched chains while also weakly associating with unbranched K48-chains. Of note, 8 of the 15 identified proteins in Cluster 3 & 4 contain annotated ubiquitin-binding domains and motifs, indicating that these may bind directly to the branched bait chains. Three of the eight proteins in Cluster 4 (ATXN3, ZFAND2B, RHBDD1) also harbor VCP/p97-binding motifs (VIM, VBM) in addition to one or more ubiquitin interacting motifs (UIMs) (**Figure 2D**).

This suggests that they may bind branched chains directly and deliver to or process branched K48-K63-Ub chains on VCP/p97. Interestingly, VCP/p97 is ranked in between ATXN3, ZFAND2B and RHBDD1, and the established VCP/p97 substrate adapters NPLOC4 and UFD1, which have been previously shown to bind K48-linked Ub chains to initiate unfolding of modified client proteins (Johnson et al., 1995; Meyer et al., 2000). Aptly, we also detect additional p97-binding proteins/substrate adapters (UBXN1, NSFL1C/p47) in Cluster 5 that contains mainly unbranched K48-linked Ub chain binders (Kondo et al., 1997; McNeill et al., 2004). These results indicate the co-existence of VCP/p97 complexes functionalied with different substrate adapters that confer a range of Ub chain preferences from unbranched K48-Ub to branched K48-K63-Ub chains (Figure 2E).

We, then attempted to validate our MS data *in vitro* using recombinant proteins for the identified branched chain specific binders. Unfortunately, for several of the interactors, we were unable to purify full-length proteins. We therefore tested if the specificity towards branched chain binding was encoded in the minimal UBD of the protein. Of the minimal UBDs tested, several of them either did not bind to the Ub chains tested or lacked specificity (**Figure S2C**). This suggests that additional regions of the protein are required to impart branched Ub binding properties. In contrast, the minimal UIM motifs of RFC1 showed high specificity at binding to branched K48-K63 chains with no detectable binding observed for the unbranched controls (**Figure 2F**). In summary, these results reveal for the first time the existence of proteins that bind specifically to branched Ub chains.

### ULTIMAT DUB assay monitors cleavage of individual linkages in Ub chains

Since we identified multiple DUBs in the pulldown with branched and unbranched Ub chains, we next investigated whether some DUBs can preferentially cleave branched Ub chains. However, conventional DUB assays monitoring the cleavage of polyUb chains lose information on which specific linkage within a Ub chain is cleaved (Pruneda and Komander, 2019). Hence, to capture the information of which Ub is being cleaved and thereby investigate DUB activity towards branched chains, we developed a precise, quantitative DUB assay, ULTIMAT (**U**biquitin **L**inkage **T**arget **I**dentification by **Ma**ss-**T**agging). The principle of the ULTIMAT DUB assay relies on the use of substrate polyUb chains in which each Ub moiety within the chain is of a specific but slightly different mass that can be distinguished by matrix-assisted laser desorption/ionization time of flight mass spectrometry (MALDI-TOF MS) (**Figure 3A**). Following incubation with a DUB, the released monoUb species are detected using MALDI-TOF MS which enables identification and quantification of the exact linkage cleaved. Employing the Ub-capping approach for stepwise chain ligation, we generated a set of four tandem mass-tagged Ub chains of unbranched (K48-Ub3, K63-Ub3) and branched architectures (2-(K63)-K48-Ub4, 2-(K48)-K63-Ub4) with individual masses incorporated in each Ub building block (**Figure 3A**). The proximal Ub of each chain is C-terminally truncated (Ub^1-72^ = 8181 Da), the central Ub is unmodified (Ub^WT^ = 8656 Da), the distal Ub harbors K48R and K63R mutations (Ub^KR^ = 8621 Da), and the second, K48-linked distal Ubs of the two branched chains are additionally ^15^N-isotope labeled (^15N^Ub^KR^ = 8730 Da). Following incubation with a DUB, the released mono-Ub species were analyzed by MALDI TOF MS and quantified relative to an internal standard of ^15^N-isotope labeled Ub (^15N^Ub = 8670 Da) (De Cesare et al., 2020) (**Figure S3A, Supplementary Data**). As controls, we first analyzed the activity of the K63-specific DUB AMSH/STAMBP and K48-specific DUB MINDY1 in this assay, which showed that they only cleaved the K63- or K48-linked Ub chains, respectively (**Figure S3B**).

**Figure 3.**
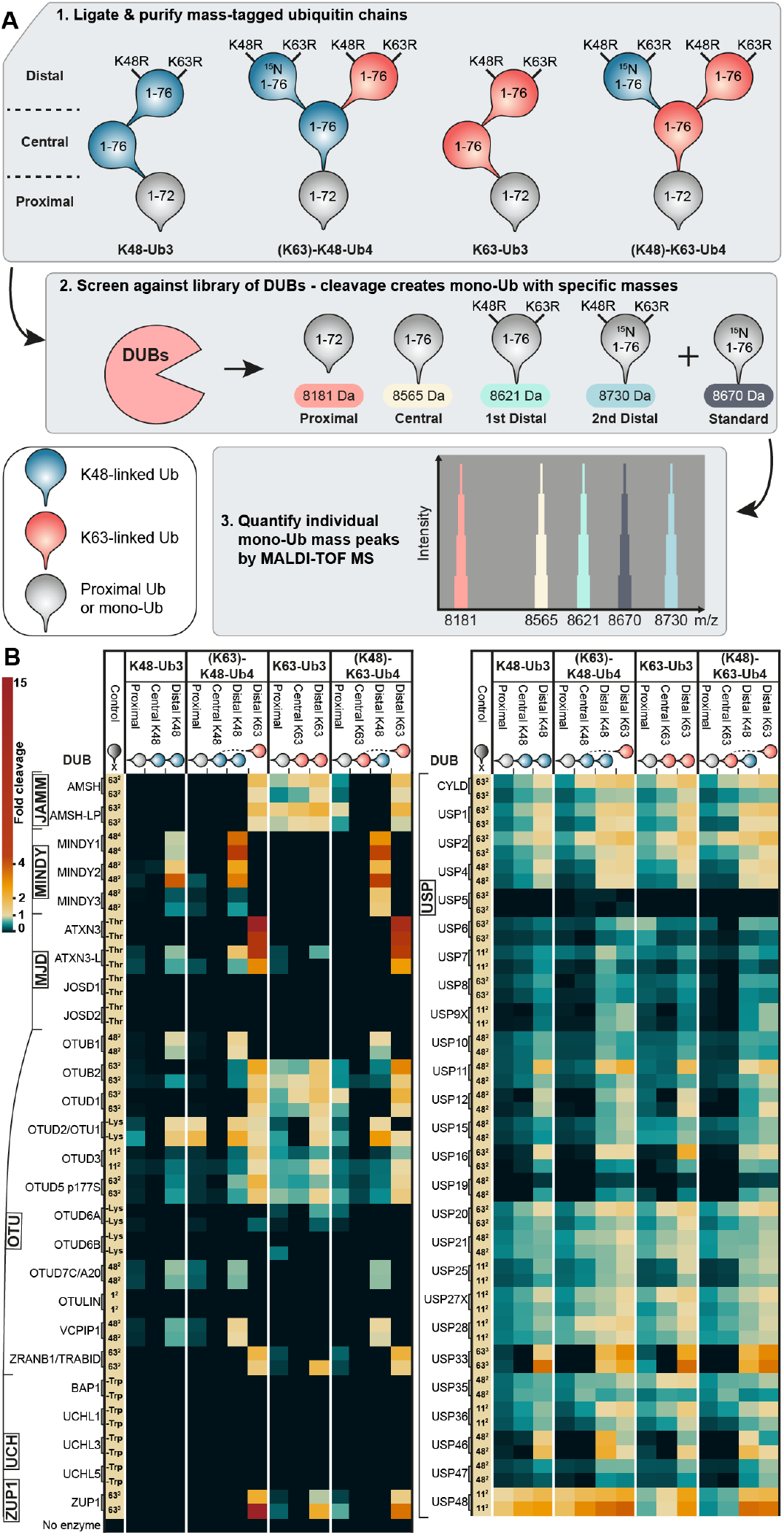
Profiling debranching activity of 53 human DUBs in ULTIMAT DUB assay. A) Principle and schematic workflow of the ULTIMAT DUB assay. B) Screen of 53 human DUBs in duplicate using ULTIMAT DUB assay with K48- and K63-linked chains. Heatmap shows individual data-points of duplicate measurements of released Ub moieties normalized to internal ^15^N-Ub standard and relative to cleaved control substrate. Schematic of substrates and location of Ub moieties are depicted above heatmap. Control substrates are either homotypic Ub chains of specific linkage type and length (e.g. 63^2^ = K63-linked diUb), Ub with C-terminal Trp residue (-Trp), Ub modified with isopeptide-linked Lys residue (-Lys), or Ub with ester-linked Thr-residue (-Thr).

Having established the robustness and reproducibility of this method, we then analyzed a panel of 53 human DUBs for their activity towards homotypic and branched Ub substrates compared to a positive control substrate (**Figure 3B**). As expected, no cleavage of K48- and K63-linked substrates were detected for the highly M1-specific DUB OTU-LIN or members of the UCH DUB family, which prefer short and disordered peptides at the C-terminus of Ub (Bett et al., 2015; Keusekotten et al., 2013). Accordingly, members of the USP family, known to be less linkage-selective, displayed a broad cleavage activity against all tested substrates, notably, with a trend to cleave from the distal end of the chain (**Figure 3B**). Of note, in our assay, we observed no inhibitory effect of the branched chain architecture on CYLD activity, as suggested in a previous study (Ohtake et al., 2016) (**Figure 3B**). Importantly, we identified some DUBs such as the MINDYs and ATXN3 that showed marked preference for cleaving branched ubiquitin chains.

### MINDY1 recognizes branched K63-linked Ub via a cryptic S1’^br^ Ub-binding site

In the ULTIMAT DUB assay, both MINDY1 and MINDY3 stood out for their high activity at cleaving K48-linkages off branched chains. The K48-specific DUB MINDY1, which usually prefers long K48-chains, cleaved the distal K48-linked Ub off the branched chains more efficiently compared to the distal Ub of unbranched K48-Ub3. MINDY1 has 5 well-characterized Ub binding sites on its catalytic domain to interact with five K48-linked Ub moieties and is virtually inactive against short chains such as K48-Ub2 (Abdul Rehman et al., 2016). However, to our surprise, MINDY1 activity was also enhanced towards 2-(K48)-K63-Ub4, a branched chain where a single K48-linked Ub branches off a K63-linked Ub3 (**Figure 3B**).

We therefore proceeded to systematically analyze processing of branched chains by all four active members of the MINDY DUB family (MINDY1-4). MINDY-1 and −2 have a similar domain architecture with a disordered N-terminus, catalytic domain, followed by tandem MIUs (motif interacting with ubiquitin) and a C-terminal CAAX box for membrane anchoring. MINDY3 consists of a catalytic domain with an inserted EF-hand feature, while MINDY4 has a large, disordered N-terminus with a C-terminal catalytic domain (**Figure 4A**). Comparing the activities of catalytic and full-length MINDYs in an ULTIMAT DUB assay with branched and unbranched K48- and K63-linked substrates (**Figure 4B**) shows that the processing of branched chains by full-length MINDY1 is accelerated 5.4-fold compared to the distal Ub of unbranched K48-Ub3, but only 2.8-fold in case of the catalytic domain, suggesting a role of for the tandem MIUs in efficient branched chain processing. In contrast, both MINDY2 full-length and catalytic domain process the distal K48-linked Ub of K48-Ub3 and the two branched Ub4 chains with similar efficiency. MINDY3 shows comparable activity against the distal Ub of unbranched K48-Ub3 and branched (K63)-K48-Ub4, but strikingly is 4.4-fold more active at cleaving the K48-linked distal Ub of (K48)-K63-Ub4. These data suggest a specific role of MINDY3 to remove K48-Ub chain linkages branching off K63-Ub chains. In contrast, MINDY4 cleaves distal K48-linkages in both unbranched K48-Ub3 and branched (K63)-K48-Ub4 with similar efficiency but displays reduced processing of (K48)-K63-Ub4 (0.5-fold). Unlike the other MINDY DUBs that are K48-linkage specific, MINDY4 also displays weak activity towards K63-linkages (Armstrong et al., in preparation). In summary, we find that each MINDY family member has a unique cleavage profile for branched K48-K63-Ub chains with MINDY1 and −3 preferring branched over unbranched substrates.

**Figure 4.**
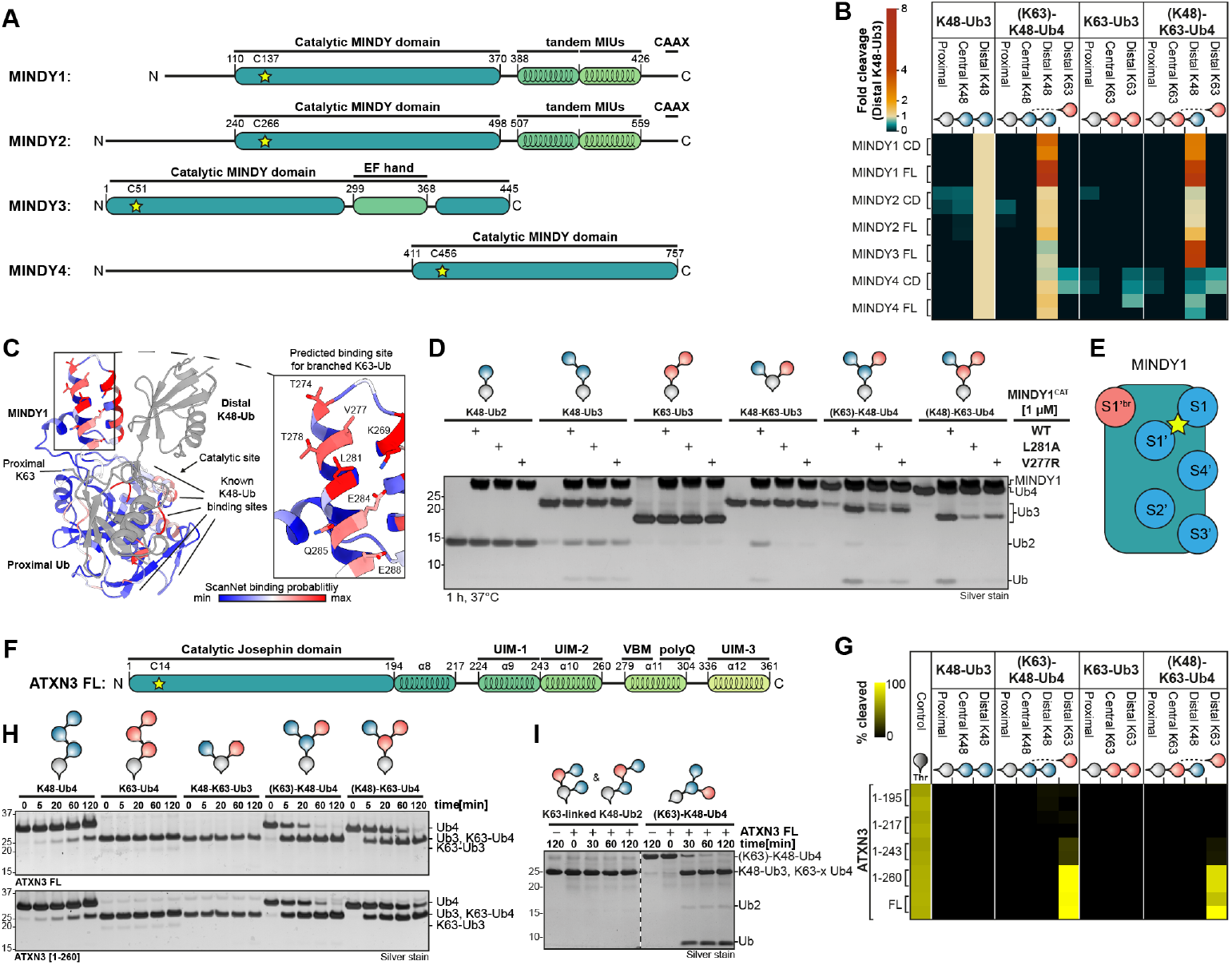
Unwinding the debranching activity of MINDYs and ATXN3. A) Schematic domain overview of active MINDY family members with highlighted catalytic Cys residues. B) ULTIMAT DUB assay in duplicate of catalytic domains and full-length constructs of MINDY family members against branched and unbranched K48- and K63-linked substrate chains. Heatmap depicts individual datapoints of duplicate measurements of released Ub moieties normalized to the internal ^15^N-Ub standard and to the intensity of the distal Ub of K48-Ub3. C) Crystal structure of the catalytic domain of MINDY1 in complex with K48-Ub2 (PDB 6TUV) with MINDY1 residues colored by ScanNet-binding probability score (blue, white, red) and Ub molecules in grey. Zoomed-in view of potential K63-Ub binding site with residues shown as stick models. D) Silver-stained SDS-PAGE analysis of DUB assays with wild-type catalytic domain of MINDY1 or point mutants in potential K63-Ub binding site (L281A or V277R) screened against a panel of branched and unbranched K48- and K63-linked Ub chains. E) Schematic of six Ub binding sites located in MINDY1’s catalytic domain colored by Ub-linkage binding preference in blue (K48) or red (K63). F) Schematic domain overview of ATXN3 with highlighted catalytic Cys residue. G) ULTIMAT DUB assay in duplicate with full-length and C-terminally truncated ATXN3 against branched and unbranched K48- and K63-linked substrate chains. Heatmap shows individual datapoints of duplicate measurements normalized to internal ^15^N-Ub standard as absolute percentage of substrate linkage cleaved. H) Silver-stained SDS-PAGE analysis of DUB assays with full-length ATXN3 (top) and ATXN3 [1-260] (bottom) against a panel of branched and unbranched K48- and K63-linked Ub chains. I) Silver-stained SDS-PAGE analysis of DUB assays with full-length ATXN3 against Ub4 chains of K63-linked Ub2 (K63-x Ub4, left) and branched (K63)-K48-Ub4.

Detailed structural and biochemical characterization of MINDY-1 and −2 have identified five Ub binding pockets for K48-linked Ub on the catalytic domains, as demonstrated by the co-crystal structure of MINDY2 in complex with K48-linked Ub5 (PDB 7NPI) (Abdul Rehman et al., 2021). However, these previously identified Ub binding sites (S1, S1’-S4’) would be unable to accommodate a K63-linked Ub of a branched K48-K63-Ub chain, as the K63 residue of the proximal Ub in the S1’-pocket is positioned opposite to these known K48-binding sites (**Figure 4C**). Strikingly, analyzing the protein binding probability of MINDY1 surface residues using the ScanNet webserver (Tubiana, Schneidman-Duhovny and Wolfson, 2022) predicted a novel high-confidence binding patch adjacent to the S1’-pocket of MINDY1 near the K63-residue of the proximal Ub (**Figure 4C**). We theorized that mutation of this potential K63-binding site in MINDY1, which is not one of the five Ub binding sites previously identified, should affect cleavage of branched K48-K63-Ub chains but not that of unbranched chains. Indeed, the introduction of either V277R or L281A into MINDY1 abolished cleavage of K48-K63-Ub3, while processing of unbranched K48-Ub3 was unaffected (**Figure 4D**). Similarly, processing of the tetrameric branched chains was severely impeded by mutations at this site, demonstrating that MINDY1’s catalytic domain has a sixth Ub binding site that can bind branched K63-linked Ub, which we refer to as S1’^br^-site (**Figure 4E**).

### ATXN3 is a debranching enzyme

In the ULTIMAT DUB assay, we also identified K63-specifc DUBs that were activated when presented with branched chain architectures. Notably, the p97-associated DUB ATXN3, which was previously thought to cleave long K63-linked chains, showed ~10-fold higher cleavage activity for the distal K63-linked Ub in both branched Ub4 substrates compared to the control Ub-Thr substrate (De Cesare et al., 2021), while no cleavage of the proximal K63-linkage or unbranched K63-Ub3 was detected (**Figure 3B**). As ATXN3 demonstrates markedly higher activity towards branched chains, we dissected this activity in further detail. ATXN3 belongs to the Josephin family of DUBs and harbors an N-terminal catalytic domain (Cat: aa 1-195) followed by a helical extension (aa 195-217), tandem UIM (UIM1: aa 224-243, UIM2: aa 243-260), a p97-interacting motif and poly-Gln repeat (VBM-polyQ: aa 279-304), and a C-terminal third UIM (UIM3: 336-361) (**Figure 4F**).

We generated truncated versions of ATXN3 to further dissect the contribution of p97 interaction and the different UBDs to its debranching activity. An ULTIMAT DUB assay comparing the different ATXN3 truncations revealed that ATXN3 with the tandem UIM (aa 1-260) is the minimal region required to efficiently cleave the branched chain architectures, while the control substrate Ub-Thr was not affected by any of the truncations (**Figure 4G**). We next performed a time course experiment to compare the activity of full-length ATXN3 and ATXN3 1-260 (**Figure 4H**). Both proteins showed weak activity at cleaving unbranched K48-Ub4 and were unable to cleave K63-Ub4 or branched K48-K63-Ub3. Strikingly, about 50% of the tetrameric branched chains containing an additional proximal Ub prior to the branchpoint were cleaved within 5 minutes by both ATXN3 constructs (**Figure 4H, S4A**).

As ATXN3 interacts with VCP/p97 via a VBM motif located between UIM2 and UIM3, we next tested if addition of VCP/p97 affects the DUB activity of ATXN3, especially since it has been suggested that VCP/p97 can activate ATXN3 ac-tivity (Laço et al., 2012) (**Figure S4B-C**). Interestingly, addition of p97 hindered cleavage of branched Ub4 substrates by both full-length ATXN3 and ATXN3^1-260^, which lacks the C-terminal VBM and UIM3 motifs. As both ATXN3 proteins are affected by VCP/p97 addition independent of the VBM motif, these results indicate that VCP/p97 competes for ubiquitylated substrates *in vitro* and thereby slows down chain cleavage by ATXN3.

It was previously reported that ATXN3 prefers cleaving long K63-linked Ub chains and K63-linkages in mixed (unbranched) Ub chains containing K48- and K63-linkages (Winborn et al., 2008). In particular, this ‘mixed’ chain was assembled by ligating wild-type K48-linked Ub2 using K63-specific E2s UBE2N/UBE2V1 (R&D Systems, personal communication, 2022), which should in theory result in a mixture of branched and mixed Ub4 chains, as one K48-Ub2 molecule could be ligated to the proximal or distal Ub moiety of the other K48-Ub2 to create branched Ub4, 1-(K48-Ub)-2-(K48-Ub)-K63-Ub2 {Ub4}, or mixed Ub4, 2-(1-(K48-Ub)-K63-Ub)-K48-Ub2 {Ub4}) (**Figure 4I**). To compare ATXN3 activity against mixed and branched chains directly, we ligated wild-type K48-Ub2 using UBE2N/UBE2V1 and purified the resulting tetra-Ub chains. A direct comparison of the cleavage of this mixed K63-linked K48-Ub2 and branched (K63)-K48-Ub4 by ATXN3 shows that only a small fraction of the mixed K63-linked K48-Ub2 is cleaved to Ub2 after 2 h, while the majority of (K63)-K48-Ub4 is trimmed to K48-Ub3 within 10 minutes (**Figure 4I**). Our results suggest that the previously observed partial cleavage of K63-linked K48-Ub2 may be attributed to a small fraction of contaminating branched chains in the K63-linked K48-Ub2 chain mixture. In summary, the ULTIMAT DUB assay presented here is a superior method for monitoring DUB activity, which reports on the precise linkages cleaved within complex polyUb and will be an invaluable approach to understand chain length, architecture and cleavage properties of DUBs. Importantly, this approach led us to identify ATXN3 as a debranching enzyme that efficiently removes K63-linkages from branched K48-K63-Ub chains.

### Development of a high-affinity nanobody against branched K48-K63-Ub chains

The absence of tools such as antibodies that specifically bind and detect branched Ub chains formed in cells is a major barrier to their facile analysis and characterization. Nanobodies are versatile tools that have been used widely in both structural biology and cell biology studies (Manglik et al., 2017). We therefore devised a screening strategy based on a synthetic yeast surface display nanobody library (McMahon et al., 2018) to obtain nanobodies that bind selectively to branched K48-K63-Ub chains (**Figure 5A**). First, we performed a negative selection step to remove yeast expressing undesired binders to unbranched K48- or K63-linked Ub chains. This was followed by positive selection to enrich for binders to branched K48-K63-Ub3 (1-(K48)-K63-Ub2 [Ub3] {1-72-AVI*biotin, K48R K63R, K48R K63R} (**Figure 5A, Figure S5A**). Following four rounds of magnetic cell sorting (MACS) selection, we identified a promising candidate nanobody, NbSL3, which bound branched K48-K63-Ub with sub-micromolar affinity (K_D_ = 740 ± 140 nM) and displayed good solubility in bacterial and mammalian cells (**Figure 5B, S5B**).

**Figure 5.**
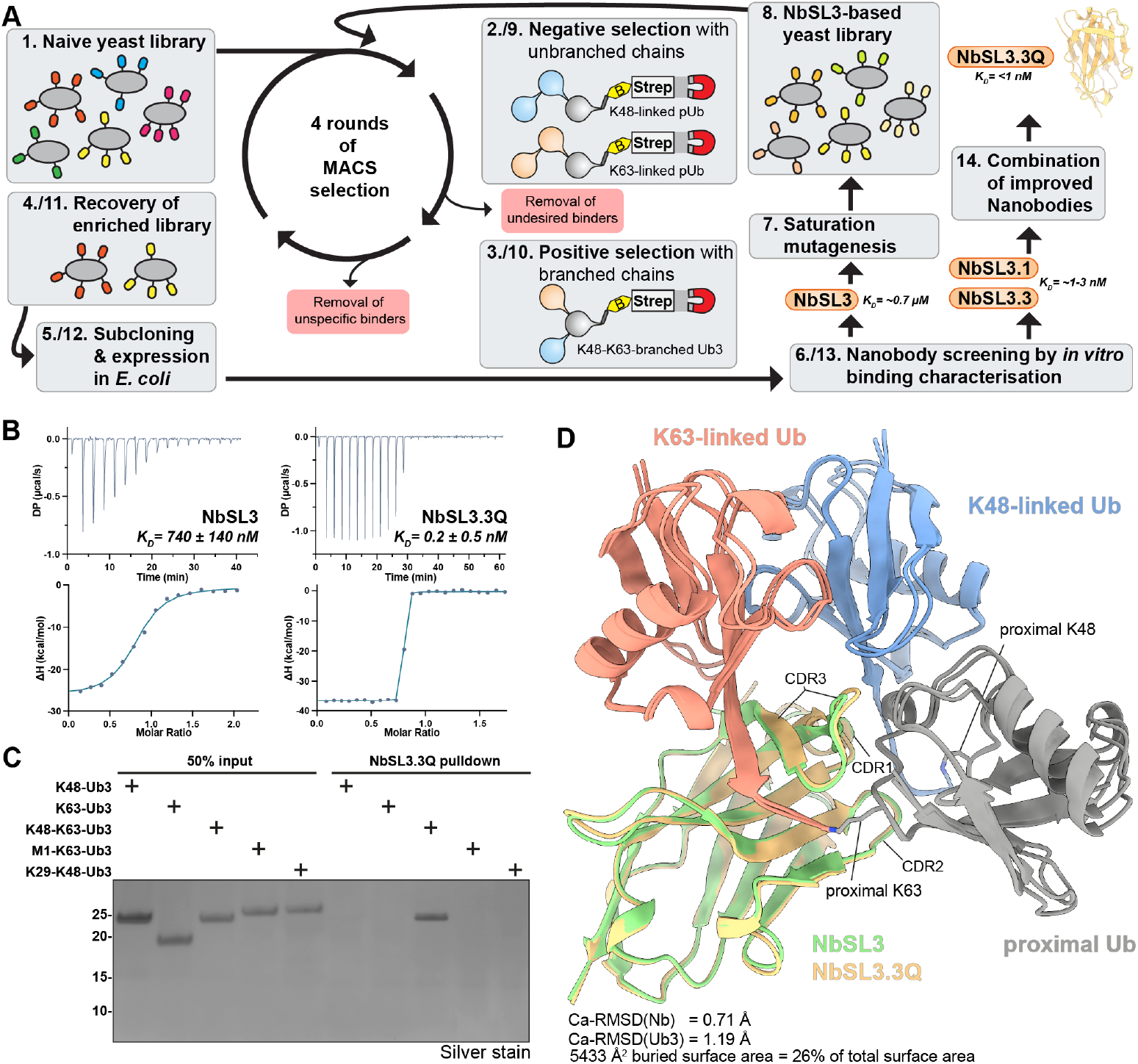
Engineering of the branched K48-K63-Ub specific, high-affinity nanobody NbSL3.3Q. A) Schematic workflow of nanobody selection and maturation using yeast-surface display screening. B) ITC analysis of first-generation nanobody NbSL3 and matured third-generation nanobody NbSL3.3Q binding to branched K48-K63-Ub3. C) Silver-stained SDS-PAGE analysis of *in vitro* pulldown with NbSL3.3Q-immoblized agarose beads against a panel of branched and unbranched Ub3 chains. D) Superimposed crystal structures of NbSL3 (green) and NbSL3.3Q (yellow) each in complex with branched K48-K63-Ub3 in cartoon representation. K48-linked Ub in blue, K63-linked Ub in red and proximal Ub in grey. Isopeptide linkages are shown as stick models.

Next, we used site-directed saturation mutagenesis to randomize individual amino acid positions in the complementarity determining regions (CDRs) of NbSL3 and generated a diverse NbSL3-based yeast library with ~2×10^8^ unique nanobody sequences for maturation of NbSL3. Following four rounds of negative and positive MACS selection, the top four nanobodies identified (NbSL3.1-4) displayed improved binding to branched K48-K63-Ub chains with affinities in the low-nanomolar range (~1-100 nM) (**Figure S5C-D**). Rational analysis of mutations in the matured nanobodies led us to combine the mutations of the top two ranking nanobodies, adding the F47Q mutation of NbSL3.1 to the three mutations of NbSL3.3 (R33V, N56V, V101E), thereby generating a third generation nanobody, NbSL3.3Q (**Figure S5C**). Excitingly, NbSL3.3Q bound to branched K48-K63 chains with pico-molar affinity (**Figure 5B**). We next analyzed the specificity by performing pulldowns using NbSL3.3Q-cross-linked to agarose resin and assayed binding to unbranched chains (K48-Ub3, K63-Ub3) and branched chains (K48-K63-Ub3, M1-K63-Ub3, K29-K48-Ub3) (**Figure 5C**). This revealed that NbSL3.3Q bound with high specificity only to branched K48-K63-linked Ub chains while no binding was detected for unbranched K48- or K63-linked chains. Additionally, NbSL3.3Q did not bind to the two unrelated branched chain types tested, K29-K48 and M1-K63, firmly establishing the specificity of this nanobody.

To understand how the nanobody specifically recognizes only branched K48-K63-Ub chains, we determined crystal structures of branched K48-K63-Ub3 in complex with the original NbSL3 or the matured NbSL3.3Q (**Figure 5D, S5E, Table S1**). In both structures, the branched Ub chain wraps around the nanobody and adopts a completely different conformation compared to the free K48-K63-Ub chain crystal structure (**Figure 1D**). Superimposition of the two structures revealed a near identical global binding mode with a slight rotation of the three Ub moieties relative to the Nb in the matured nanobody structure (Ca-RMSD(Nb) = 0.71 Å, Ca-RMSD(Ub3) = 1.19 Å). In both structures, the C-terminal residues V70, L71 and L73 of the two distal Ub moieties engage in hydrophobic interactions with the nanobody’s complementarity-determining regions (CDRs) and the proximal Ub. CDR1 interacts with the C-terminus of the K48-linked Ub and the proximal Ub, CDR2 makes contacts with both K48- and K63-linkages on the proximal Ub, and the large CDR3 loop inserts itself deeply in between all three Ub moieties and directly contacts the K63-linkage (**Figure 5D**). A PISA analysis (Krissinel, 2015) of the NbSL3.3Q complex revealed a buried surface area of ~5430 Å^2^, which corresponds to 26% of the total surface area, indicating a compact complex. We performed a comparative analysis of the original NbSL3 and matured NbSL3.3Q structures to establish the roles of the four mutations of NbSL3.3Q (R33V, F47Q, N56V, V101E) in improving the binding affinity by almost three orders of magnitude. To our surprise, only V101E contributed to a *de novo* interaction, while the other three mutations instead appeared to reduce steric hinderance and relieved unfavorable contacts of the non-matured nanobody, thereby promoting tighter binding (**Figure S5E**). Overall, the extensive interactions with Ub and the direct recognition of the K48- and K63-linkages provides a structural basis for the high affinity and specificity observed towards branched K48-K63-Ub chains. This direct mode of branched chain recognition differs from that of the branched K11-K48-Ub chain antibody which works as a co-incidence detector to recognize the presence of both K11 and K48-linked Ub (Yau et al., 2017).

### Application of nanobodies to investigate cellular functions of branched chains

Having developed a selective high affinity nanobody, we performed a pulldown to analyze cellular proteins that are modified with branched K48-K63-Ub chains. As a positive control, we treated U2OS cells with the proteasome inhibitor MG-132, as branched K48-K63-Ub chains were previously linked to proteasomal degradation (Ohtake et al., 2018). As expected, we observed an accumulation of high molecular weight ubiquitylated species (HMW-Ub) in U2OS cell extracts following proteasome inhibition, but, to our surprise, we detected no enrichment of branched K48-K63-Ub chains in subsequent pulldowns with NbSL3.3Q (**Figure 6A**). As we identified multiple VCP/p97-binding proteins to be associated with branched K48-K63-Ub chains and the p97-associated DUB ATXN3 to be highly active at cleaving branched chains, we wondered if branched K48-K63-Ub chains serve as signals for VCP/p97-mediated processes. To test this hypothesis, we treated U2OS cells with the allosteric, small molecule VCP/p97 inhibitor NMS-873, the pro-teasomal inhibitor MG-132, the HSP70 inhibitor VER-155008 and the N-glycosylation inhibitor Tunicamycin to induce unfolded protein response (Azim and Surani, 1979; Ito et al., 1975; Lee and Goldberg, 1998; Magnaghi et al., 2013; Williamson et al., 2009). Excitingly, while both p97 and proteasomal inhibition led to a significant accumulation of HMW-Ub conjugates, only the ubiquitylated proteins stabilized upon p97 inhibition were captured by the branched K48-K63-Ub specific nanobody NbSL3.3Q (**Figure 6A**). Similarly, inhibition of p97 with the ATP-competitive inhibitor CB-5083 also led to significant accumulation of branched K48-K63-Ub chains (**Figure 6B**). These results suggest that proteins modified with branched K48-K63-Ub chains are clients for processing by VCP/p97.

**Figure 6.**
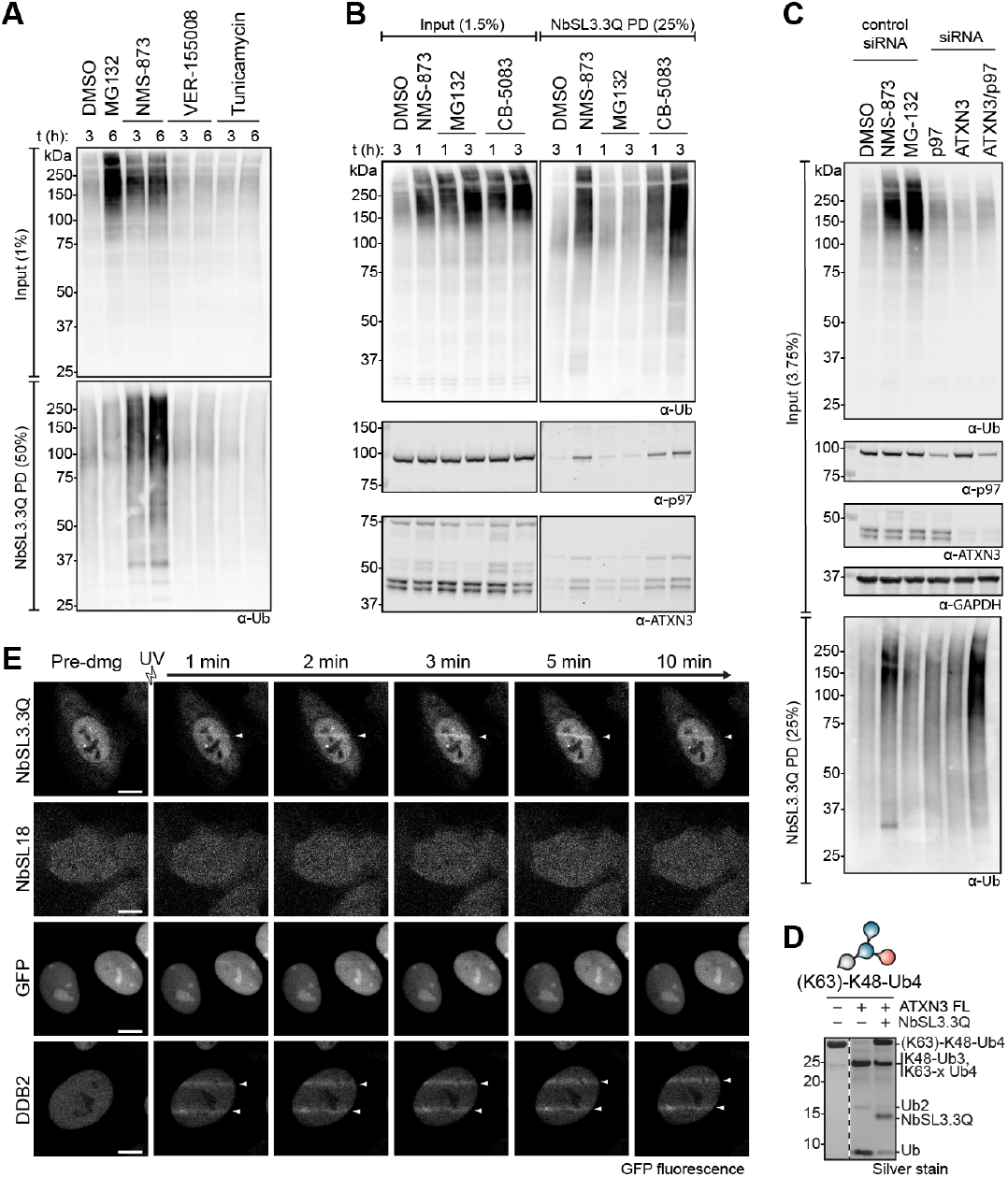
Branched K48-K63-Ub chains are made in response to VCP/p97 inhibition and at sites of DNA damage. A-C) Pulldowns using NbSL3.3Q-immobilized agarose from U2OS cells A) treated with indicated inhibitors (NMS-873 − 5 *μ*M, MG-132 − 10 *μ*M, VER-155008 − 5 *μ*M, tunicamycin − 5 *μ*g/ml). Western blot analysis of total ubiquitin in input lysate and eluted proteins. B) treated with VCP/p97-inhibitors NMS-873 (5 *μ*), CB-5083 (5 *μ*M) and proteasomal inhibitor MG132 (10 *μ*M). Western blot analysis of total Ub, VCP/p97 and ATXN3. C) treated with VCP/p97-inhibitors NMS-873 (5 *μ*M), CB-5083 (5 *μ*M) and proteasomal inhibitor MG132 (10 *μ*M) and non-specific siRNA or siRNA targeting p97 or ATXN or both in combination. Input and eluted proteins analyzed by Western blot against total Ub, p97, ATXN3 and GAPDH. D) Silver-stained SDS-PAGE of DUB assay with full-length ATXN3 and (K63)-K48-Ub4 incubated at 30 °C for 2 h following addition of branched K48-K63-Ub specific nanobody NbSL3.3Q. E) Live cell images of UV-micro irradiation assay with U2OS cells expressing either NbSL3.3Q-GFP, NbSL18-GFP, GFP or GFP-DDB2. Cells were imaged pre-damage and over a time course of 10 min following insult by UV laser striping.

Intriguingly, pulldowns of the branched K48-K63 ubiquitylated proteins that accumulate upon p97 inhibition also co-precipitate p97 and ATXN3 (**Figure 6B**). To further investigate the interplay between p97 and the debranching DUB ATXN3 in processing branched K48-K63-Ub chains, we performed transient siRNA knockdowns of p97, ATXN3 or co-depletion of both in U2OS cells and analyzed the formation of branched Ub (**Fig 6C**). While individual knock-down of p97 and ATXN3 did not result in an accumulation of high-MW Ub species, the combined depletion of p97 and ATNX3 led to a significant accumulation of proteins modified with branched K48-K63-Ub. Taken together, these results strongly suggest branched K48-K63-Ub chains to be signals for p97 that are regulated by ATXN3.

To further cement these observations, we monitored if branched K48-K63-Ub chains are formed on Ub(G76V)-GFP, a ubiquitin fusion degradation (UFD) reporter substrate that requires p97 activity for its unfolding and subsequent degradation (Beskow et al., 2009). Treatment of HEK293 cell lines expressing Ub(G76V)-GFP with p97 inhibitors leads to marked stabilization of the reporter (**Figure S6A**). Strikingly, a pulldown with NbSL3.3Q revealed that this p97 substrate is indeed modified with branched K48-K63-Ub chains.

The observation that p97 inhibition is required to stabilize branched K48-K63-Ub chains in cells suggests that K48-K63-Ub signals are transient and modified proteins are rapidly processed. We imagined that one potential strategy to stabilize the short-lived branched Ub chains in cells would be to use the branched K48-K63-Ub specific nanobody NbSL3.3Q to inhibit recognition and cleavage of these chains by DUBs. To test this possibility, we monitored the debranching activity of ATXN3 following addition of NbSL3.3Q to an *in vitro* DUB cleavage assay with (K63)-K48-Ub4. Indeed, the addition of equimolar amounts of NbSL3.3Q had a strong inhibitory effect, resulting in markedly reduced cleavage of the branched chain by ATXN3 in the presence of the nanobody (**Figure 6D**). This suggests that expression of NbSL3.3Q in cells would indeed stabilize and enrich cellular proteins modified with branched chains. We therefore generated cell lines for inducible expression of C-terminally GFP-tagged NbSL3.3Q (NbSL3.3Q-GFP) and performed GFP pulldowns. Indeed, immunoprecipitation of NbSL3.3Q-GFP showed enrichment of branched chains both in untreated and p97-inhibitor treated cells (**Figure S6B**). We then treated the captured HMW-Ub chains with the K48-specific DUB Miy2/Ypl191c, the K63-specific DUB AMSH, the non-specific DUB USP2, and the K63-specific de-branching enzyme ATXN3. Removal of K48-linkages with Miy2 treatment led to a reduction in the intensity of the polyUb smear and a shift towards lower MW ubiquitylated bands, and to complete removal of K48 linkages (**Figure S6C**). On the other hand, treatment with AMSH led to a reduction in total Ub signal intensity, but not to a lower MW shift and, as expected, K48-linkages were unaffected by AMSH treatment. Similarly, treatment with ATXN3, which is highly active against K63-linkages in branched K48-K63-Ub, led to a substantial decrease of total Ub intensity, but not to a shift in MW of the total HMW-Ub signal nor K48-linkage intensity. As a positive control, incubation with the non-specific DUB USP2 abolished all ubiquitin chain signals (**Figure S6C**). Together these findings suggest that the architecture of branched K48-K63-Ub chains formed in response to p97 inhibition consists predominantly of K63-linked Ub branches decorating K48-linked Ub chain trunks.

As NbSL3.3Q-GFP is expressed uniformly in cells with distribution in both cytoplasmic and nuclear compartments, we wanted to use it to track branched chain formation and localization in live cells. A critical function of VCP/p97 is the extraction of stalled transcription machineries from sites of DNA damage (Davis et al., 2012; Meerang et al., 2011). To test whether the branched K48-K63-Ub is induced by DNA damage and formed at sites of damage, we used NbSL3.3Q-GFP expressing U2OS cells in a micro-irradiation assay to induce localized DNA damage by striping with a UV laser. Live cell imaging of the irradiated cells revealed NbSL3.3Q-GFP to be rapidly recruited to sites of DNA damage within 1-2 min and was sustained over a period of at least 10 min (**Figure 6E, S6D**). The positive control GFP-DDB2 (DNA damage-binding protein 2) is also recruited to sites of DNA damage whereas GFP on its own or the unrelated GFP-tagged nanobody NbSL18 did not show recruitment to UV-laser stripes. In summary, these results reveal formation of branched Ub chains following DNA damage and in p97-related processes.

## Discussion

Branched Ub chains are increasingly being appreciated as unique signals used for information transfer in cells. They have been implicated in NF-κB signaling, cell cycle control, ERAD and protein quality control pathways. However, a better understanding of the roles of branched chains and how they function as unique signals is hampered by a lack of tools and innovative methods to systematically decipher these complex chains. A primary task is to identify molecular players that assemble, decode and erase branched chains. Barring the bispecific antibody that was developed to detect branched K11-K48-Ub chains, the field relies on the overexpression of ubiquitin variants in cells or sophisticated mass-spectrometry based approaches that are low throughput and not widely tractable (French et al., 2021). As demonstrated here, we have circumvented these issues by developing a comprehensive approach to assemble complex Ub chain architectures *in vitro* for the development of specific detection tools and identification of cellular interactors, to ultimately reveal how a particular branched chain transmits information in the cell.

### Decoding K48-K63 branched Ub signals

Compared to homotypic (unbranched) chains, branching creates unique interfaces that can be exploited by UBDs and DUBs to achieve selective recognition. We here suggest that it will be important to consider branched tetraUb chains as this creates an additional unique interface compared to commonly used branched triUb. Indeed, our results suggest that binding proteins and DUBs can discriminate between the type of branch formed on tetraUb, i.e., K48-linked Ub branching off a K63-linked Ub chain, or vice versa. Additionally, DUBs can also differentiate between branched triUb and tetraUb. Hence, for each branched type it will be important to consider the linkage composition of the ‘trunk’ of the chain as they encode distinct information. While this increases the complexity of branched chains, the methods outlined here make their analysis straightforward.

A long-standing question in the field is whether cellular proteins capable of specifically binding branched Ub chains and discriminating these from unbranched chains exist. By developing an approach to generate and immobilize polyUb chains of defined architectures such that the branched interfaces are available for protein interactions, we here provide the first instance of branched chain-specific binders. Importantly, our findings provide a handle to investigate branched Ub function. Interestingly, not much is known about several of the branched chain-specific binders identified here, thus opening new research avenues to study cellular processes regulated by branched chains. The identification of RFC1 and MORC3 suggest roles in regulating chromosome replication, replication stress, antiviral responses and interferon signaling. It is intriguing that several kinases such as PRKCZ, ROCK2 and RIOK3 are also identified as branched K48-K63-Ub chain-associated proteins. Whether these kinases are activated upon polyUb binding analogous to the TAK1 kinase (Wang et al., 2001) will be an interesting question to address. The identification of p97-related proteins, the HSP70 co-chaperone DNAJB2 and the ER-associated degradation (ERAD) associated protein RHBDD1 (Fleig et al., 2012) also suggest roles for branched K48-K63 chains in protein quality control. Importantly, we identify the p97-associated proteins ZFAND2B, RHBDD1 and ATXN3 to bind selectively to K48-K63-branched chains (**Figure 2D**), that suggest roles for branched K48-K63 chains as signals for VCP/p97. Investigating how branched Ub recognition is achieved by these proteins is likely to reveal novel mechanisms employed for polyUb recognition. The very existence of specific readers to branched chains highlights not only distinct roles but also the high precision of the ubiquitin system.

### Branched ubiquitin signals for VCP/p97

In addition to identifying p97-associated proteins as branched K48-K63 binders, we find the p97-associated DUB ATXN3 to de-branch K48-K63-Ub chains. Further, we found p97 inhibition with small molecules and depletion of p97 and ATXN3 via RNA interference to lead to an accumulation of branched K48-K63 chains in cells, and a model substrate that relies on p97 for unfolding and degradation is modified with branched K48-K63-Ub chains. Taken together, these findings strongly imply that branched K48-K63-Ub chains are a signal for VCP/p97-related processes. The wide distribution of VCP/p97 and VCP/p97-associated adapters across pulldowns with branched K48-K63 and unbranched K48-linked Ub chains suggests that these adapters can provide specialization to a variety of p97-complexes to recognize and process substrates modified with distinct Ub signals (Alexandru et al., 2008; Raman et al., 2015). We speculate that these VCP/p97-complexes might carry additional factors for quality control or fate determination post p97-processing and might help the cell to enable prioritization of processing based on the detailed architecture of Ub signals. In fact, how p97 recognizes different polyUb architectures is poorly understood and very little is also known about the role of the polyUb binding preferences of p97 adapters in substrate recognition and processing (Ahlstedt et al., 2022). Interestingly, a recent study demonstrated a role for p97 adapters in modulating the efficiency of ubiquitin-dependent substrate unfolding by p97 (Fujisawa et al., 2022). It is a distinct possibility that other p97 adapters specifically bind different branched chain types, such as branched K11-K48-Ub (Yau et al., 2017), a prospect that will be important to explore systematically.

Branched K11-K48-Ub chains have been shown to be efficient signals to trigger degradation by the proteasome partly due to their increased affinity for the proteasome receptor RPN1 over unbranched chains (Boughton et al., 2020; Meyer and Rape, 2014). While branched K11-K48 chains were associated with VCP/p97 via the adapters FAF1, p47 and UBXD7 (Yau et al., 2017), we find a different set of VCP/p97 adapters to bind branched K48-K63 chains. Moreover, we find the abundance of K48-K63 chains to not increase following proteasome inhibition and only following p97 inhibition. These observations suggest that branched chains of specific architectures may be formed under specific conditions to trigger recruitment of VCP/p97.

### Degradative vs non-degradative functions

A significant proportion of K63-linked chains in cells have been found to be branched with K48 linkages making them an abundant modification in cells (Ohtake et al., 2016; Swatek et al., 2019). Such branching could influence outcome in two ways. In the first scenario, branching of K48-linked Ub off K63-linked Ub chains could protect the chain from cleavage by DUBs while retaining the ability of the K63-linked portion of the chain to signal. Such a mechanism has been shown for NF-κB where the ligase HUWE1 branches the K63 chains by binding via its UBA domain to catalyze addition of K48 linkages (Ohtake et al., 2016). Our identification of proteins such as kinases, RFC1 and MORC3 as branched K48-K63-Ub chain binders would further support non-degradative signaling functions for these branched chains. Alternatively, branching of K63-Ub chains by the addition of K48 linkages could rapidly convert the modification to a degradative signal, thereby terminating K63 signaling by degrading the branched K48-K63 ubiquitylated signaling node. Previous work has identified K48-K63-branched chains to be abundantly present at the proteasome (Ohtake et al., 2018). We here have identified VCP/p97 and protein quality control-associated proteins to bind to branched K48-K63-Ub chains which may support a degradative role for these branched architectures. The enforced degradation of proteins using PROTAC methods is an emerging technology with widespread interest and recent work has identified PROTAC targets to be modified with branched chains of different architectures (Akizuki et al., 2022). It is imaginable that the formation of branched structures enables degradation of these otherwise stable and difficult to degrade proteins by targeting them to VCP/p97 to unfold them first. Hence, the outcome of branched K48-K63-Ub modification could either be degradative or non-degradative and would be context dependent. Our findings and the tools presented here will enable further research to explore these possibilities.

### Debranching enzymes

Our discovery of the VCP/p97-associated DUB ATXN3 as a debranching enzyme was made possible using the ULTI-MAT DUB assay we pioneered here. Interestingly, the only other debranching enzyme known to date is the pro-teasome-associated DUB UCHL5 (also referred to as UCH37) (Deol et al., 2020). The debranching activity of UCHL5 requires association of the DUB with the proteasome subunit RPN13 and the hydrophobic patches on both distal Ub moieties of a branched Ub3 chain (Du et al., 2022; Song et al., 2021). In contrast to UCHL5 that can debranch Ub3 chains, ATXN3 activity requires longer branched Ub4 substrates. Importantly, previous studies that claim ATXN3 to cleave long polyUb chains, only observe moderate activity after several hours. In contrast, the rapid processing of branched chains in under 5 minutes demonstrates a preference for branched Ub chains by ATXN3. Further structural studies are required to establish how the branched architecture is recognized by ATXN3 and how the branch point is positioned across the catalytic site. Also, whether ATXN3 is specific for K48-K63-branched Ub4 or if it could also debranch branched chains containing other linkages remains to be explored in future research.

The fact that the two main molecular machines responsible for unfolding and degradation, VCP/p97 and the proteasome, are associated with a debranching enzyme suggests that debranching is an essential prerequisite for further substrate processing. Indeed, the importance of VCP/p97-ATXN3 interaction is highlighted by VCP/p97 mutations in the proteinopathy disorder inclusion body myopathy with Paget disease of bone and frontotemporal dementia (IBMPFD). Intriguingly, degenerative disease-causing mutations in VCP/p97 stabilize and greatly enhance its interaction with ATXN3, suggesting an inhibitory role (Custer et al., 2010; Fernández-Sáiz and Buchberger, 2010; Rao et al., 2017). In contrast, loss of ATXN3 also impairs ERAD and protein degradation (Zhong and Pittman, 2006). Taken together with our findings, we propose that while branched chains are effective signals for substrate recognition by VCP/p97, ATXN3 serves an essential role at VCP/p97 to debranch the bifurcated architectures for unfolding and degradation.

A previous study identified a significant proportion of the K63-Ub chains made during NF-κB signaling to be branched with K48-linkages and such branching was attributed to enhanced NF-κB signaling as the K63-linked portion of the chains are now protected from the DUB CYLD that is unable to cleave these chains (Ohtake et al., 2016). However, we do not find CYLD activity to be affected by chain branching *in vitro*, but instead observe CYLD to have a slight preference to cleave distal Ub and K63-linked over K48-linked Ub.

An intriguing finding from the screen of 53 human DUBs using the ULTIMAT assay and branched substrates is that branched chains were preferentially cleaved by DUBs that were previously thought to prefer long, homotypic chains (Abdul Rehman et al., 2016; Kwasna et al., 2018; Winborn et al., 2008). This observation demonstrates that DUB cleavage specificity and activity data generated from homotypic Ub chains may need to be reevaluated with a panel of heterotypic chains. It also raises the question if any DUBs that were thought to be inactive, based on activity assays with homotypic chains, may have evolved to specifically and efficiently cleave branched architectures.

A frequently encountered difficulty hindering our understanding of the biological functions of DUBs is a high degree of redundancy within the ubiquitin system, limiting the effectiveness of classical knock-out or knock-down approaches. The combination of the methods presented here that permit connecting Ub chain architectures to cellular function and identifying the enzymes involved in processing these chains with the UTLIMAT DUB assay, may therefore offer an invaluable accompanying strategy. The UTLIMAT DUB assay reveals that each of the four active MINDY family members, whose biological role remains elusive to date, displays a unique activity profile when encountering branched K48-K63-Ub chains. Two members, MINDY-1 and −3, show ~5-fold increased activity against branched K48-Ub compared to homotypic chains and MINDY-3 also appears to be specifically activated by K48-Ub branching off a K63-linked trunk. With the establishment of a link between branched K48-K63-Ub chains and VCP/p97, this may now spark future research into the function of MINDYs in these processes.

Several studies have found concomitant increases of K48- and K63-linked polyUb in processes including DNA repair, NF-κB signaling, proteotoxic stress (Baranes-Bachar et al., 2018; Emmerich et al., 2013; Ohtake et al., 2016). For instance, the findings that both K48- and K63-linked chains are formed following DNA damage (Baranes-Bachar et al., 2018; Meerang et al., 2011; Panier and Durocher, 2009; Singh et al., 2019) prompted us to explore whether these are present within branched chains and led us to identify branched K48-K63 chains at sites of DNA damage (**Figure 6E**). We therefore suggest that reexamining previous findings using the new tools and methods presented here is likely to uncover hitherto unappreciated roles for branched K48-K63 chains in regulating cellular homeostasis.

We envision that the methods presented here for generating complex and functionalized ubiquitin chains *in vitro* will be applied to study other branched chain architectures in the future. Similarly, the ULTIMAT DUB assay provides a quantitative, high-throughput method to monitor cleavage of complex Ub substrates, which is an important improvement compared to existing methods that either provide only qualitative information or utilize fluorescent tags that occupy a large surface area of Ub and may thereby affect cleavage activity.

### Limitations of Study

A key feature of branched Ub chains is the unique three-dimensional architecture produced, and it is important to note that the nanobody we have developed here recognizes this architecture. Hence, one limitation of this nanobody is that it cannot be used as a detection tool in immunoblotting presumably because the unique three-dimensional epitope recognized by the nanobody isn’t preserved. On the other hand, the nanobody confers the advantage of being able to detect branched chain formation in living cells as it can be expressed in human cells. Further, the ability of the Nb to bind to and protect branched Ub chains from cleavage in cells could be leveraged to study branched chain function. Importantly, the Nb can be used to investigate branched chain formation under native conditions that does not involve expression of Ub variants.

Despite having identified DUBs with debranching activity, one limitation of the ULTIMAT DUB assay is that it uses Ubs with Lys-to-Arg mutations which may in some instances affect DUB activity. We attempted to minimize the effect of these mutations on our results by including the same Lys-to-Arg mutations in the unbranched substrates used. This is exemplified by USP5, which does not show activity against K63-linked chains bearing Lys-to-Arg mutations in the ULTIMAT DUB assay but is active against K63-linked chains assembled from wild-type Ub (**Figure 3B, S3C**). Indeed, in an existing USP5~Ub structure (PDB 3HIP), both K48 and K63 residues of the distal Ub are tightly engaged in the S1 pocket of USP5, explaining the inhibitory effect of the Lys to Arg mutations (**Figure S3D**). Similarly, we cannot rule out if the Lys-to-Arg mutations present in Ub chains used in pulldowns from cell lysate may have prevented binding to certain proteins and thereby their identification.

The comprehensive approach we employed resulted in identifying K48-K63 branched chains as a signal in VCP/p97-related processes. Future work will establish if dedicated adapter-VCP/p97 complexes recognize K48-K63 branched chains, what cellular stress conditions trigger the modification of substrates with K48-K63 branched chains and how these substrates are processed by VCP/p97. While our work identifies ATXN3 as a debranching enzyme, it is not known how debranching by ATXN3 fine tunes substrate recognition and processing by VCP/p97. Further, the present study has only examined ATXN3 activity towards K48-K63 branched chains and whether it can debranch other branched architectures needs to be examined. Structural studies are also needed to determine how branched chains are recognized by ATXN3.

## Acknowledgements

We thank members of the Kulathu lab for helpful discussion and critical reading of the manuscript, especially Logesvaran Krshnan. We thank Yoshua Kristariyanto for the HEK293 Flp-In Trex Ub(G76V)-GFP stable cell line. We thank Andrew Kruse (Harvard Medical School) for making the synthetic yeast surface display library available to the wider scientific community and Ignacio Moraga Gonzalez (University of Dundee) for helpful advice during nanobody selection. We thank Mateusz Gregorczyk & Thomas Carroll (MRC PPU) for assistance with UV-laser stripe live cell imaging and image processing. This work was supported by funding from ERC Consolidator grant (Y.K. grant 101002428), MRC grant MC_UU_00018/3, the Lister Institute of Preventive Medicine (Y.K). This paper was typeset with the bioRxiv word template by @Chrelli: www.github.com/chrelli/bioRxiv-word-template.

## Author contributions

Y.K. and S.M.L. conceptualized the study. S.M.L., D.K., L.S., I.C., L.A.A., A.K., and C.J. expressed and purified proteins. S.M.L and M.R.M. cloned expression plasmids. S.M.L. prepared and purified ubiquitin chains, nanobody reagents and cross-linked agarose beads. S.M.L, I.W., and I.C. performed nanobody selection, maturation, and characterization. S.M.L., M.R.M., Y.K. and F.L. planned, executed, and analyzed MS pulldown experiments. S.M.L, V.D.C., and Y.K. developed, conducted, and analyzed ULTIMAT DUB assays. S.M.L. and L. A.A. ran gel-based DUB assays. M.R.M. generated and maintained cell cultures, concluded cell-based experiments and Western Blotting. M.R.M. and S.M.L. completed and analyzed UV-stripe assay. S.M.L. and Y.K. wrote the manuscript with input from other authors.

## Experimental Procedures

### Materials availability

All cDNA constructs in this study were generated by S.M.L, M. R.M. and the cloning team at the MRC PPU Reagents & Services. All plasmids have been deposited with the MRC PPU Reagents and Services and are available at https://mrcppureagents.dundee.ac.uk/.

**Table 1.**
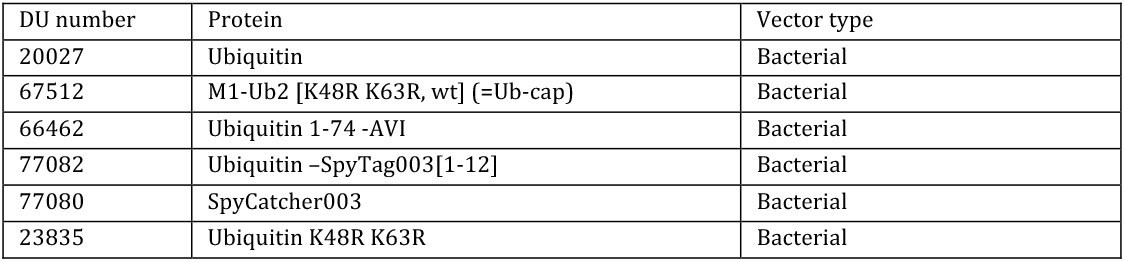
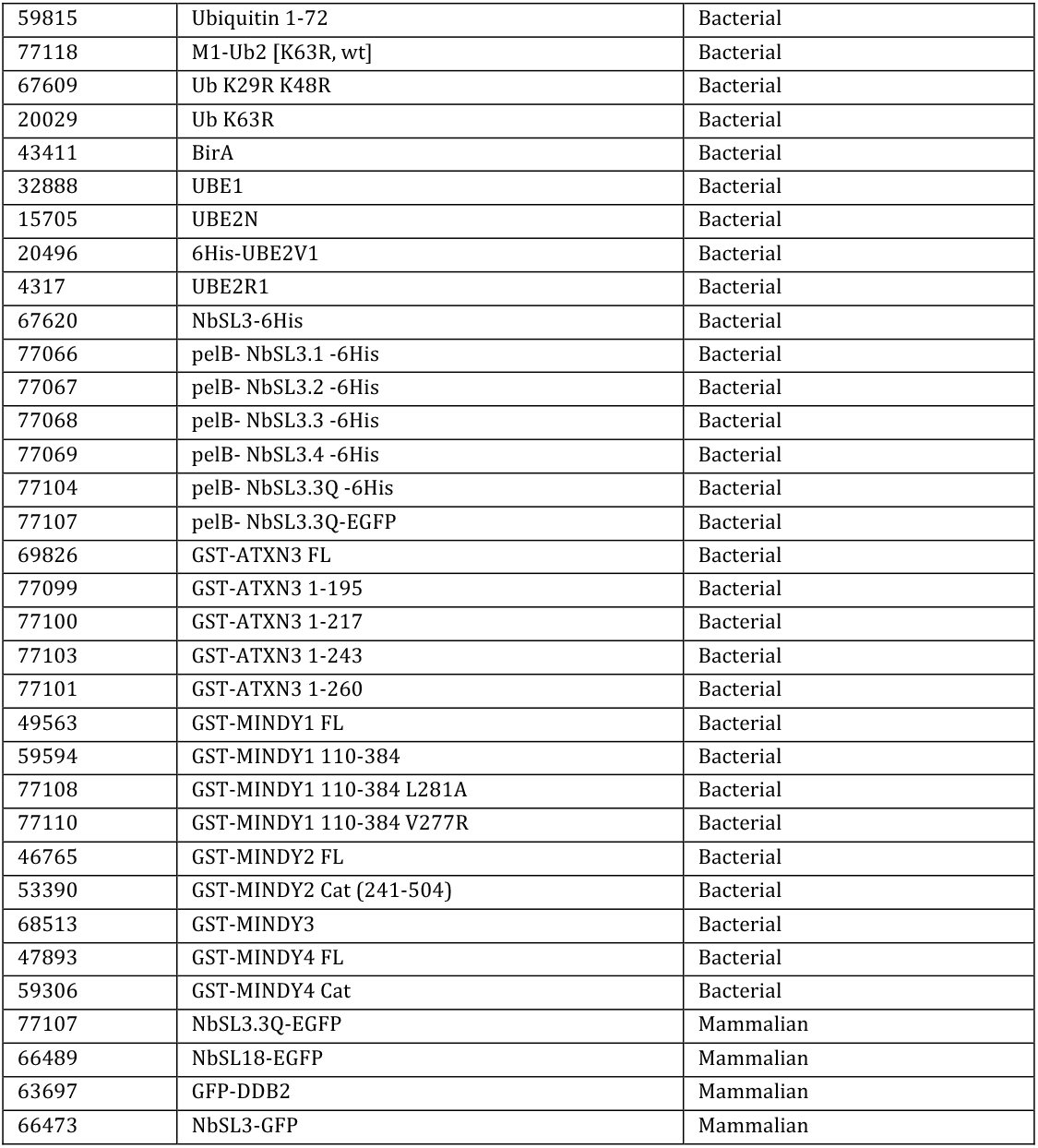
List of plasmids.

| DU number | Protein | Vector type |
| --- | --- | --- |
| 20027 | Ubiquitin | Bacterial |
| 67512 | M1-Ub2 [K48R K63R, wt] (=Ub-cap) | Bacterial |
| 66462 | Ubiquitin 1-74 -AVI | Bacterial |
| 77082 | Ubiquitin -SpyTag003[1-12] | Bacterial |
| 77080 | SpyCatcher003 | Bacterial |
| 23835 | Ubiquitin K48R K63R | Bacterial |
| 59815 | Ubiquitin 1-72 | Bacterial |
| 77118 | M1-Ub2 [K63R, wt] | Bacterial |
| 67609 | Ub K29R K48R | Bacterial |
| 20029 | Ub K63R | Bacterial |
| 43411 | BirA | Bacterial |
| 32888 | UBE1 | Bacterial |
| 15705 | UBE2N | Bacterial |
| 20496 | 6His-UBE2V1 | Bacterial |
| 4317 | UBE2R1 | Bacterial |
| 67620 | NbSL3-6His | Bacterial |
| 77066 | pelB- NbSL3.1 -6His | Bacterial |
| 77067 | pelB- NbSL3.2 -6His | Bacterial |
| 77068 | pelB- NbSL3.3 -6His | Bacterial |
| 77069 | pelB- NbSL3.4 -6His | Bacterial |
| 77104 | pelB- NbSL3.3Q -6His | Bacterial |
| 77107 | pelB- NbSL3.3Q-EGFP | Bacterial |
| 69826 | GST-ATXN3 FL | Bacterial |
| 77099 | GST-ATXN3 1-195 | Bacterial |
| 77100 | GST-ATXN3 1-217 | Bacterial |
| 77103 | GST-ATXN3 1-243 | Bacterial |
| 77101 | GST-ATXN3 1-260 | Bacterial |
| 49563 | GST-MINDY1 FL | Bacterial |
| 59594 | GST-MINDY1 110-384 | Bacterial |
| 77108 | GST-MINDY1 110-384 L281A | Bacterial |
| 77110 | GST-MINDY1 110-384 V277R | Bacterial |
| 46765 | GST-MINDY2 FL | Bacterial |
| 53390 | GST-MINDY2 Cat (241-504) | Bacterial |
| 68513 | GST-MINDY3 | Bacterial |
| 47893 | GST-MINDY4 FL | Bacterial |
| 59306 | GST-MINDY4 Cat | Bacterial |
| 77107 | NbSL3.3Q-EGFP | Mammalian |
| 66489 | NbSL18-EGFP | Mammalian |
| 63697 | GFP-DDB2 | Mammalian |
| 66473 | NbSL3-GFP | Mammalian |

### Protein expression

Recombinant proteins were expressed in E. coli BL21(DE3) in Autoinduction medium containing 100 μg/ml Ampicillin or 50 μg/ml Kanamycin, as appropriate, at 18-25°C for 24 h at 180 rpm shaking speed. Cells were harvested by centrifugation at 4000 g for 20 min at 4°C. To prepare isotope-labelled ^15^N-ubiquitin, E. coli were grown in ^15^N-minimal medium (8 g/L glucose, 2 g/L ^15^NH4Cl2, 1 x M9 salts, 2 mM MgSO4, 0.2x Studier trace metals, 1x MEM vitamins) supplemented with 50 μg/ml Kanamycin to OD_600_ 1.5 at 37 °C and expression was induced with 1 mM IPTG for 20 h at 20 °C. For wild-type and mutant ubiquitin, cells were resuspended in 20 ml Ub lysis buffer (1 mM EDTA, 1 mM AEBSF, 1 mM Benzamidine) and lysed by sonication. The pH of the lysate was adjusted by addition of 100 mM sodium acetate pH 4.5 and incubated for 3-16 h at 20 °C. The lysate was adjusted to 50 mM sodium acetate through addition of water prior to clarification by centrifugation at 30k g and 4 °C for 20 min. Ubiquitin was purified by ion exchange chromatography on a Resource S column (6 ml) in 50 mM sodium acetate pH 4.5 using a NaCl salt gradient. The pH of elution fractions was adjusted by addition of 100 mM Tris-HCl pH 8.5 prior to concentration in 3 kDa MWCO centrifugal filter units (Amicon) and finally buffer exchanged into 50 mM Tris-HCl pH 7.5.

For purification of cytoplasmic proteins, pellets from 1 liter of expression culture were resuspended in 20 ml bacterial lysis buffer (50 mM Tris-HCl pH 7.5, 300 mM NaCl, 0.5 mM TCEP, 1 mM Benzamidine, 1 mM AEBSF) and lysed by sonication. Lysates were clarified by centrifugation at 30k g and 4 °C for 30 min and applied to affinity resin for subsequent purification. GST-tags were removed by overnight incubation with 3C-protease at 4 °C. For crystallization, protein complexes were purified by gel filtration (Superdex 200 pg 16/600) equilibrated in 20 mM HEPES pH 7.5, 150 mM NaCl.

For periplasmic proteins, cells from 1 liter expression culture were resuspended in 20 ml high-osmotic lysis buffer (50 mM Tris-HCl pH 7.5, 150 mM NaCl, 20% sucrose, 1 mM EDTA, 1 mM Benzamidine, 1 mM AEBSF and 5 mg hen egg lysozyme) and incubated for 20 min at 20 °Ç. The cell suspension was centrifuged at 15k g and 4 °C for 10 min and pellet and supernatant were separated. The pellet was resuspended in low-osmotic lysis buffer (50 mM Tris-HCl pH 7.5, 1 mM EDTA, 1 mM Benzamidine, 1 mM AEBSF) and incubated on a roller at 4 °C for 40 min. The high-osmotic supernatant and 5 mM MgCl2 were added to the low-osmotic cell suspension and the mixture was centrifuged at 30k g at 4 °C for 20 min. The supernatant containing released periplasmic proteins was subjected to affinity purification.

### Ubiquitin chain ligation, purification, and modification

Ubiquitin chains were assembled from 1.5 mM ubiquitin in 40 mM Tris-HCl pH 7.5, 10 mM MgCl, 10 mM ATP at 30 °C for 2-16 h. Formation of K48-linkages was catalyzed by 1 μM UBE1 and 25 μM UBE2R1 and K63-linkages by 1 μM UBE1, 20 μM UBE2N, 20 μM UBE2V1. Ub chains were separated by length using ion exchange chromatography on a Resource S column (6 ml) in 50 mM sodium acetate pH 4.5 using a NaCl salt gradient. The pH of elution fractions was adjusted by addition of 100 mM Tris-HCl pH 8.5 prior to concentration in 10 kDa MWCO centrifugal filter units (Amicon) and chains buffer exchanged into 50 mM Tris-HCl pH 7.5. Biotinylation of 200 μM AVI-tagged ubiquitin chains was catalyzed by addition of 1 μM BirA in 50 mM Tris-HCl pH 7.5, 5 mM MgCl2, 2 mM ATP and 600 μM biotin for 2 h at 25 °C. Subsequently, the protein was buffer exchanged into 50 mM Tris-HCl pH 7.5 to remove free biotin. Successful biotinylation was assessed through a Streptavidin-shift assay by incubating biotinylated protein with 5-fold excess streptavidin for 5 min at 20 °C, addition of 1x LDS sample buffer and subsequent SDS-PAGE analysis., where the stable strep-tavidin-biotin complex induces a ~60 kDa MW-shift.

### Immobilization of proteins on agarose beads

Proteins were coupled to amine-reactive NHS-activated agarose resin (Abcam, ab270546) based on the manufacturer’s protocol. Briefly, the protein was buffer exchanged into Coupling buffer (50 mM HEPES pH 7.5, 500 mM NaCl). Per 1 mg of protein, 1 ml of NHS-activated resin was activated by washing with 50 ml ice-cold Acid buffer (1 mM HCl), then quickly equilibrated by washing with 50 ml icecold Coupling buffer and mixed with the protein. The coupling reaction was allowed to proceed on an end-over-end roller at 4 °C for 16 h. After coupling, the resin was washed six times total, alternating between 50 ml High pH buffer (0.1 M Tris-HCl pH 8.5) and 50 ml Low pH buffer (0.1 M sodium acetate pH 4.5, 0.5 M NaCl), to remove any non-covalently bound protein. Finally, the resin was equilibrated with Storage buffer (50 mM Tris-HCl pH 7.5, 150 mM NaCl, 0.02% sodium azide) as a 50% slurry and stored at 4 °C. SpyTag-ubiquitin chains were incubated with SpyCatcher-coupled agarose beads for 16 h at 20 °C on an end-over-end roller, washed six times with High pH and Low pH buffer as described above, and stored as a 50% slurry in Storage buffer.

### Nanobody selection & maturation

Specific nanobodies against K48-K63-Ub3 were selected from a naïve yeast display library, generously shared by the Kruse lab (McMahon et al., 2018) and yeast culture and magnetic cell sorting (MACS) was performed as previously described (Armstrong et al., 2021). Briefly, yeast were cultivated in YGLC-glu medium (80 mM sodium citrate pH 4.5 (Sigma), 6.7 g/l yeast nitrogen base w/o amino acids (BD Biosciences), 2% glucose, 3.8 g/l Do mix-trp) at 30 °C and 200 rpm shaking speed for 16 h. Nanobody expression was induced by growth in YGLC-gal (same as YGLC-glu, but glucose replaced with galactose) at 20 °C and 200 rpm for 48-72 h. Nanobodies were selected in four round of MACS by negative selection against 400 nM biotinylated, homotypic K48- and K63-linked polyUb and positive selection with decreasing concentrations (2000 nM to 400 nM) of biotinylated, branched K48-K63-Ub3. Following MACS, the total DNA of yeast colonies grown on YGLC-glu agar plates was isolated by resuspending single colonies in 100 μl 200 mM lithium acetate and 1% SDS, followed by incubation at 70 °C for 5 min and brief vortexing after adding 300 μl ethanol. The mixture was centrifuged at 15k g for 3 min and the pellet washed once with 70% ethanol before resuspension in 100 μl H2O. Following additional centrifugation at 15k g for 1 min to remove cell debris, the supernatant was transferred to a fresh microtube and 1 μl was used as template DNA for a 25 μl PCR reaction (KOD HotStart, Millipore) to amplify the Nanobody insert using primers NbLib-fwd-i (CAGCTGCAGGAAAGCGGCGG) and NbLib-rev-i (GCTGCTCACGGTCACCTGG). Nanobodies were subcloned into pET28a vectors for periplasmic expression in bacteria with an N-terminal pelB signal sequence and C-terminal 6His-tag.

NbSL3 was matured through directed evolution of the nanobody binding properties by construction of a NbSL3-based maturation library using saturation mutagenesis. Two-step multiple overlap extension PCR (MOE-PCR) was performed based on the procedure described by McMahon et al. (2018) to generate a DNA library encoding ~1.97 * 10^8 NbSL3 variants each harboring up to 4 mutations in either of the variable positions of CDR1, CDR2, CDR3 loops or additional two residues of NbSL3, Thr75 and Tyr77, in a fourth loop located between beta-sheets β7 and β8, we refer to as CDR2.5. The codons of the variable amino acid positions in these four regions were replaced with degenerate NNK codons, which encode all 20 natural amino acids and only a single stop codon (see Table 2). For MOE-PCR with KOD HotStart polymerase, equimolar primer pools encoding each CDR region (NbSL3_P3, −P5, −P7 and P9) were used to prepare a 10 μM NbSL3 primer mix combining all ten NbSL3-encoding primers (NbSL3_P1 - P10). A 5-fold dilution series of 2 μl primer mix was used in 25 μl MOE-PCR assembly reactions in 15 cycles of denaturation (20 s, 95 °C), annealing (20 s, 60 °C) and elongation (10 s, 70 °C), followed by 15 cycles of amplification after addition of 0.3 μM flanking primers pYDS_fwd_1 and pYDS_rev_1 with an increased annealing temperature of 68 °C. The Nb insert DNA band of 462 bp size was purified from a 2% agarose gel and served as a template for two subsequent PCR amplifications rounds using the primer pairs pYDS_fwd_2 + pYDS_rev_2 and pYDS_fwd_3 + pYDS_rev_2 to generate matching overhangs for homologous recombination with the yeast surface display vector pYDS649. Electroporation of yeast with the NbSL3 DNA library was performed following the protocol developed by Benatuil and coworkers (Benatuil et al., 2010), Briefly, a 100 ml culture of the yeast strain BJ5465 was grown to OD600 of 0.4 and co-transformed by electroporation with 24 μg the amplified NbSL3 DNA library and 8 μg linearized pYDS digested with BamHI and NheI. Highly efficient electroporation was achieved on a BTX 630 Exponential Decay Wave Electroporation System (Harvard Bioscience, Inc.) set at 2500 V, 200 Ω and 25 μF resulting in time constants between 3-4 ms. A dilution series of transformed yeast was streaked out on YGLC-glu agarose plates to estimate a transformation efficiency of >95%. The transformed yeast library was recovered in 500 ml YGLC-glu selection medium and used in 4 rounds of MACS as described above, but with K48-K63-Ub3 concentrations decreasing from 400 nM to 100 nM. Following maturation, Nb sequences in individual yeast colonies were sequenced and subcloned into pET28a vector for bacterial expression and subsequent characterization.

**Table 2.**
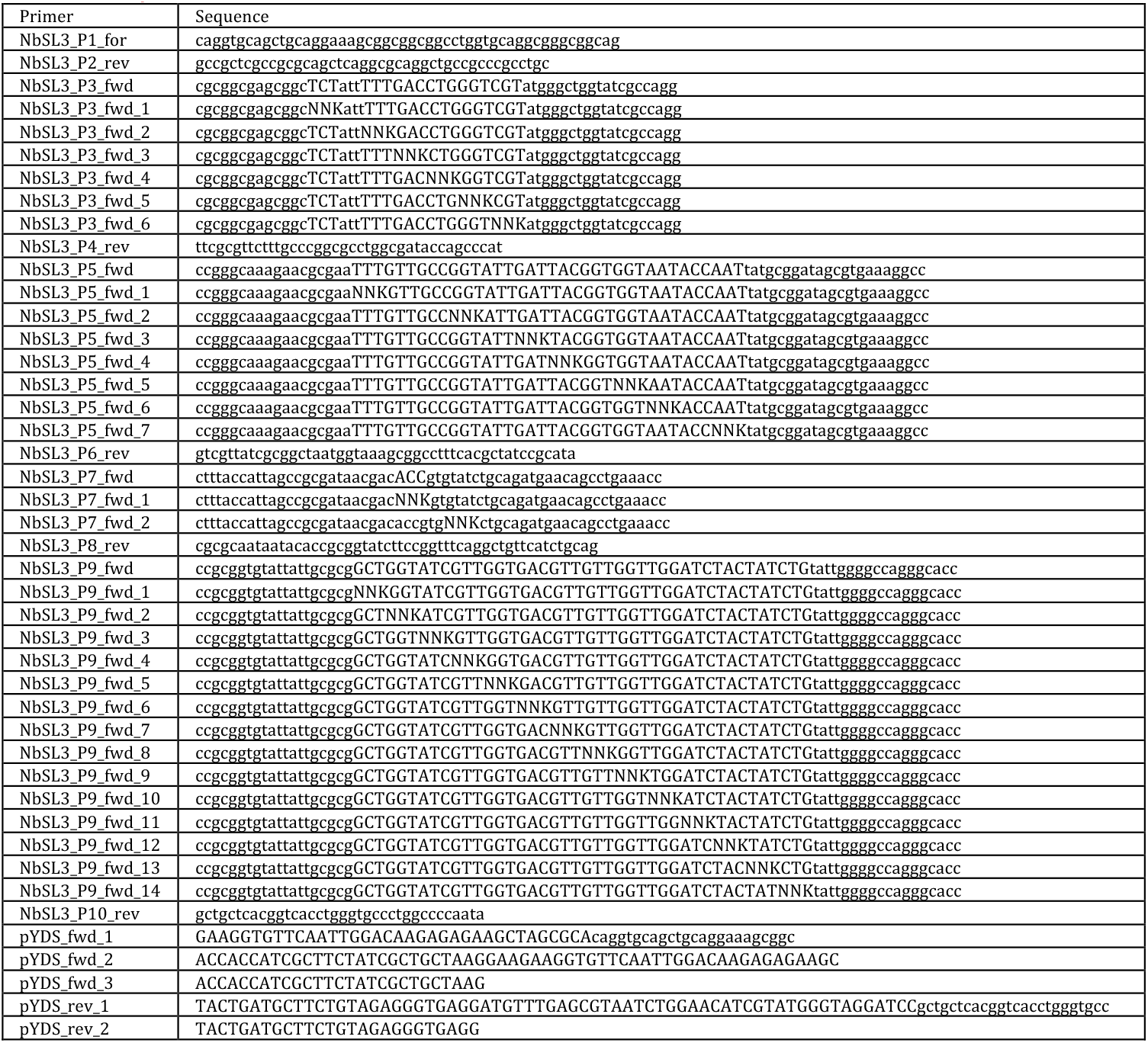
List of primers used for NbSL3 maturation.

### Isothermal titration calorimetry (ITC)

ITC measurements were executed at 25 °C on a MicroCal PEAQ-ITC instrument (Malvern). Immediately prior to analysis, proteins were dialyzed into degassed ITC buffer (20 mM HEPES pH 7.5, 150 mM NaCl) at 4 °C for 16 h. The data was analyzed with MicroCal Analysis Software (Malvern) and fitted using a one-sided binding model to calculate binding constants.

### Protein crystallization, data collection and structure determination

All protein crystals were obtained by sitting drop vapor diffusion method. K48-K63-Ub3 crystals were obtained at 22 mg/ml in 0.2 M Ammonium acetate, 20 mM Tris pH 7.5, 50 mM NaCl, 0.1 M Sodium citrate tribasic dihydrate pH 5.6, 30% w/v PEG 4000 at 20 °C. Crystals harvested and cryoprotected with Mother liquor supplemented with 30% glycerol. The complex of NbSL3 and K48-K63-Ub3 was crystallized at 12 mg/ml in 0.1 M Bis-Tris pH 7.2, 0.28 M MgCl2, 21% PEG3350, 0.15 M NaCl, 0.05 M Tris/HCl pH 7.5 at 4 °C. Crystals harvested and cryo-protected with Mother liquor supplemented with 30% glycerol. The complex of the matured NbSL3.3Q and K48-K63-Ub3 was concentrated to 14.5 mg/ml in 20 mM HEPES pH 7.5, 150 mM NaCl. Mixed 200 nl protein with 100 nl mother liquor (0.1 M HEPES pH 7.5, 10% 2-propanol, 20% PEG4000). Crystals harvested and cryo-protected with Mother liquor supplemented with 30% glycerol. All datasets were collected at the ESRF beamline ID23-2 and solved by molecular replacement with ubiquitin (PDB ID 1UBQ) or the nanobody scaffold of Nb.b201 (PDB ID 5VNW). Detailed data collection and refinement statistics are documented in the Supplementary information.

### Gel-based deubiquitination assays

DUBs were incubated in DUB buffer (50 mM Tris-HCl pH 7.5, 50 mM NaCl, 10 mM DTT) at 20 °C for 10 min to fully reduce the catalytic Cys. Deubiquitination assays were typically performed with 1 μM DUB and 2.5 μM substrate Ub chain in DUB buffer at 30 °C, unless stated otherwise. Reactions were stopped by addition of 1x LDS sample buffer and cleavage of Ub chains analyzed by SDS-PAGE and silver staining using the Pierce Silver stain kit (Thermo Fisher) according to the manufacturer’s instructions but skipping the initial wash step in water to avoid wash-out of monoUb.

### ULTIMAT DUB assay

Sample preparation, spotting on MALDI target and MALDI-TOF MS analysis were performed as previously described [De Cesare 2020, 2021]. Briefly, DUBs and substrates were diluted in the reaction buffer (40 mM Tris-HCl pH 7.5, 1 mM TCEP and 0.01% BSA). 3 μl of recombinantly expressed DUBs were aliquoted in 384 Eppendorf Lowbind well plates. Control ubiquitin chains (M1, K11, K48, K63 dimers, Ub-Threonine, Ub-Lysine, Ub-Tryptophan, K63 trimer and K48 tetramer), ULTIMAT ubiquitin trimers and branched chains were separately added to each reaction at the final concentration of 1.2 μM. Reaction buffer was used to bring the total volume reaction to 10 μl. Samples were incubated at 30° for 30 minutes. The reaction was stopped with 2.5 μl of 6% TFA supplemented with 4 μM ^15^N ubiquitin (to be used as internal standard). 384 plates were centrifuged at 4000 rpm for 3 minutes. Spotting on 1536 AnchorChip MALDI-TOF target was performed in technical duplicate using a 5 decks Mosquito nanoliter pipetting system. Samples were analyzed using a Rapiflex MALDI-TOF instrument equipped with Compass for FlexSeries 2.0 and flexControl version 4.0 Build 48 software version in Reflectron Positive mode. Detection window was set between 7820 and 9200 m/z. Movement on sample spot was set on Smart - complete sample, allowing 4000 shots at raster spot within an 800 μm diameter. Acquired spectra were automatically integrated using FAMS FlexAnalysis method (version 4.0 - Build 14), SNAP Peak detection algorithm, SNAP average composition Averagine, Signal to Noise Threshold of 5, Baseline Subtraction TopHat. SavitzkyGolay algorithm was used for smoothing processing. ^15^N ubiquitin signal (“heavy” ubiquitin, 8669.5 m/z) was used to internally calibrate each data point. Spectra were further manually verified to ensure mass accuracy throughout the automated run. Peak areas of interest were exported into csv file and manually analyzed using Microsoft Excel. Average peak areas of released monoUb resulting from the cleavage of substrates, i.e., ubiquitin control chains (8565.7 m/z) or ULTIMAT branched chains (8181.3; 8622.2; 8729.9 and 8565.7 m/z), were independently normalized to the internal ^15^N-Ub standard (8669.5 m/z) and quantified using the following equation:

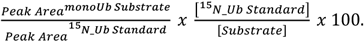

Datasets were normalized to the individual control substrates of each DUB (DUB panel, Figure 3B) or to the intensity of the distal K48-Ub of the K48-Ub3 substrate (MINDY panel, Figure 4B). Data were visualized in python using the Plotly graphing library (Plotly Technologies Inc., 2015).

### Cell culture

U2OS, U2OS Flp-In Trex, and HEK293 Flp-In Trex cell lines were maintained in DMEM (Gibco) supplemented with 10% FBS (Gibco), 2 mM L-glutamine (Gibco), and 100 U/ml Penicillin/Streptomycin (Gibco), and incubated at 37°C with 5% CO2 unless otherwise stated. Trypsin(0.05%)-EDTA (Gibco) was used to dissociate cells for passage. All cell lines were routinely tested for mycoplasma.

### Stable cell line generation

For generation of cell lines stably expressing tetracycline-inducible GFP-tagged constructs, Flp-In Trex cells were co-transfected with a 1:9 ratio (w/w) of GFP vector:pOG44 Flp recombinase vector using PEI Max 40k (Polysciences). To select for integrant cells, 24 hours post-transfection, media was switched out for fresh DMEM supplemented with 200 μg/ml hygromycin B. Selection media was periodically refreshed, and cultures monitored until all mock transfected control cells were dead. Tet-inducible expression of the proteins of interest were subsequently confirmed via Western blotting with an anti-GFP antibody, following overnight incubation with 1 μg/ml tetracycline (Figure S6B). In experimental use, the NbSL3.3Q-GFP construct was induced with 0.1 μg/ml tetracycline, with 1 μg/ml tetracycline used for all others.

### Chemicals and compounds

Cell culture treatments were carried out using the following chemicals at the indicated concentrations: DMSO (Sigma), MG-132 (Sigma) - 10 μM, NMS-873 (Sigma) - 5 μM, tunicamycin (abcam) - 5 μg/ml, VER-155008 (Sigma) - 10 μM, CB-5083 (Generon) - 5 μM, tetracycline hydrochloride (Sigma) - 0.1-1 μg/ml, BrdU (Sigma)-10 μM.

### RNA interference

RNAi was carried out using Lipofectamine RNAiMAX (Thermo Scientific) according to the manufacturer’s protocol. Briefly, cells were seeded into 6-well plates at 1-2×10^5^cells/well. The following day, cells were transfected with 25 pmol of siRNA duplexes prepared in RNAiMAX reagent. Cells were then incubated at 37 °C for 48 hours prior to harvest and subsequent analysis. RNA sequences used are presented in table 3.

**Table 3.**
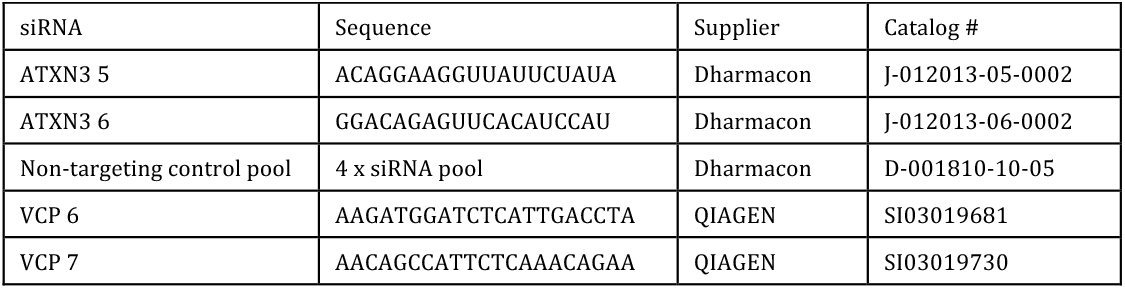
siRNA sequences used in this study.

### Pulldown with HALO-tagged UBDs and recombinant Ub chains

HALO-tag fusion constructs of UBDs were used for pulldown with recombinant Ub chains as previously described (Kristariyanto et al., 2017). Briefly, 10 nmol of HALO-tagged UBDs were immobilized on 100 μl HALOLink resin (Promega) in 500 μl HALO-coupling buffer (50 mM Tris pH 7.5, 150 mM NaCl, 0.05% NP-40 substitute, 0.5 mM TCEP) rolling at 4 °C for 2 h. Beads were spun at 800 g for 2 min to remove supernatant, washed 3 times with HALO-wash buffer (50 mM Tris pH 7.5, 250 mM NaCl, 0.2% NP-40, 0.5 mM TCEP) and resuspended in 100 μl ice-cold HALO-pulldown buffer (50 mM Tris pH 7.5, 150 mM NaCl, 0.1% NP-40, 0.5 mM TCEP, 0.5 mg/ml BSA). Per pulldown, 20 μl of coupled HALO-resin (50% slurry) were added to 30 pmol chains in 480 μl HALO-pulldown buffer and incubated at 4 °C turning end-over-end for 1 h. Beads were spun at 800 g and 4 °C for 2 min and washed twice with 500 μl HALO-wash buffer and transferred to a fresh 1.5 ml microtube for the final wash with 500 μl HALO-coupling buffer. Each pulldown was resuspended in 20 μl 1.33x LDS sample buffer and analyzed by SDS-PAGE and silver stain.

### Pulldown with NbSL3.3Q-coupled agarose beads and recombinant Ub chains

Recombinant Ub chains were diluted to 1 μM in NbSL3.3Q-pulldown buffer (20 mM HEPES pH7.5, 150 mM NaCl, 0.5 mM EDTA, 0.5% NP-40) and 2.5 μg of each chain was used per pulldown. 20 μl of agarose beads coupled with 1 mg/ml NbSL3.3Q and pre-equilibrated in NbSL3.3Q-pulldown buffer was incubated with Ub chains on a roller at 4 °C for 1 h. Beads were pelleted by spin at 500 g and 4 °C for 2 min and washed 5 times with ice-cold NbSL3.3Q-pulldown buffer. Washed beads were resuspended in 20 μl 2X LDS sample buffer and analyzed by SDS-PAGE and silver stain.

### Pulldown with NbSL3.3Q-coupled agarose beads and cell lysate

Cells were lysed in Co-IP (50 mM Tris-HCl pH 7.5, 150 mM NaCl, 0.5 mM EDTA, 0.5% NP-40) or RIPA (Thermo Scientific) lysis buffers supplemented with 1X cOmplete Protease Inhibitor (Roche), 1 mM AEBSF (Apollo Scientific), 5 mM chloroacetamide (Sigma), and 0.02% Benzonase (Sigma). Following clarification, protein content of lysates was assessed via Bradford assay (Thermo Scientific), and samples diluted to 0.5-2 mg/ml in Co-IP lysis buffer. Samples were mixed with 10 μl of NbSL3.3Q-coupled agarose beads per 500 μg of cell lysate and incubated on a roller at 4 °C for 1 hour. Beads were washed 4 times with Co-IP lysis buffer, and proteins eluted in 2X LDS sample buffer. Elution fractions were separated from beads by applying to SpinX filter columns and spun at 2,500 x g for 2 minutes. Input and elution fractions were subsequently analyzed by SDS-PAGE followed by immunoblotting.

### GFP-pulldown

For GFP pulldown, lysates were prepared using Co-IP lysis buffer as above, then mixed with 10 μl of GFP binder-agarose beads (MRC PPU Reagents and Services) per 500 μg of cell lysate and incubated at 4°C on a roller for 1 hour. Beads were washed, and proteins eluted and subsequently analyzed as described above.

### Western blotting

Protein samples were mixed with 4X LDS Sample buffer and 10X Reducing Agent (both Thermo Scientific) and incubated at 70 °C for 10 minutes. Following SDS-PAGE and protein transfer, membranes were stained with Ponceau S (Sigma) to assess loading and transfer efficiency. If intended for ubiquitin blotting, membranes were boiled in milliQ water for 10 minutes prior to blocking to ensure denaturation of ubiquitin chains. Chemiluminescent blots were subsequently visualized via a ChemiDoc MP (BioRad) using Clarity or ClarityMAX ECL reagents (BioRad), and fluorescent blots with an Odyssey Clx (LiCor Biosciences).

### Antibodies

Antibodies were sourced from the indicated manufacturers and used at 1:2000 dilution unless otherwise stated: anti-GFP (abcam, ab290), anti-VCP/p97 (Proteintech, 10736-1-AP, 1:4000), anti-ATXN3 (Proteintech, 13505-1-AP), anti-Ubiquitin (Biolegend, P4D1), anti-GAPDH (Proteintech, 10494-1-AP, 1:5000). Secondary detection was carried out using anti-rabbit or anti-mouse HRP-conjugated (CST, 7074, 7076, both 1:5000) or IRDye800CW/680RD-conjugated (LiCor Biosciences, 926-32211, 926-32210, 926-68073, 926-68070, all 1:15000) antibodies.

### MS Pulldown with SpyTag-Ub chains immobilized on Spy-Catcher-agarose beads

For each pulldown, 25 μg of SpyTag-Ub chain were immobilized on 50 μl SpyCatcher agarose beads (1 mg SpyCatcher cross-linked per 1 ml of NHS-activated agarose) by incubation in a total volume of 150 μl 50 mM HEPES pH 7.0 at 22 °C for 16 h while gently rotating end-over-end. The beads were spun down at 500 g for 2 min and washed three times with SpyTtag-Wash buffer (10 mM Tris pH 7.5, 150 mM NaCl, 0.1 mM EDTA, 1x complete protease inhibitor (Roche)) and resuspended as a 50% slurry in wash buffer. Sixteen 15 cm dishes of U2OS cells were grown to ~90% confluency in DMEM + 2 mM L-glutamine + 100 U/ml PenStrep + 1 mM Na pyruvate + 10% FBS at 37°C in 5% CO2 atmosphere and each washed with 5 ml PBS before harvesting by scraping cells into 1 ml ice-cold lysis buffer (10 mM Tris pH 7.5, 150 mM NaCl, 0.5 mM EDTA, 50 mM NaF, 1 mM NaVO4, 0.5% NP-40, 1x complete protease inhibitor, 0.02% benzonase, 1 mM AEBSF, 1 mM NEM) per dish. Lysates were flash frozen in liquid nitrogen and stored at −80 until further use. Per pulldown, 1 mg of lysate was incubated with 25 μg of immobilized Ub chains for 2 h at 4 °C gently rotating end-over-end. Resin was pelleted at 500 g and 4 °C for 2 min and washed 4 times with SpyTag-Wash buffer. Bound proteins were eluted by addition of 50 μl 10% SDS in 100 mM TEAB and incubation for 10 min on ice followed by centrifugation in Spin-X centrifuge tube filters at 8k g for 1 min. Samples were reduced by addition 10 mM TCEP pH 7.0 and incubation at 60 °C for 30 min shaking at 1000 rpm. Samples were cooled to 23 °C before alkylation with 40 mM iodacetamide for 30 min shaking at 1000 rpm in the dark. Samples were acidified with 1.2% phosphoric acid and diluted with 7 volumes of S-trap buffer (90% MeOH, 100 mM TEAB). Samples were loaded on S-trap mini columns and centrifuged at 1000 g and 23 °C for 1 min. The columns were washed 4 times with 400 μl S-trap buffer and transferred to a clean 2 ml tube. Per column, 10 μg trypsin (Pierce Trypsin Protease, MS Grade - Thermo Fisher) freshly dissolved in 100 μl 100 mM TEAB was added and columns briefly centrifuged at 200 g and 23 °C for 1 min. The flow-through was reapplied to the column, columns capped and incubated at 37 °C for 16 h without shaking. Peptides were eluted from columns in three steps: 80 μl 50 mM TEAB and centrifuge at 1000 g for 1 min, 80 μl 0.15% formic acid and centrifuge at 1000 g for 1 min, 80 μl 50% Acetonitrile + 0.2 % formic acid and centrifuge at 1000 g for 1 min. Combined elutions were frozen at −80 °C and freeze dried in a SpeedVac centrifuge.

### LC-MS/MS data collection

The peptides were resuspended in 0.1% formic acid in water and 2 μg loaded onto a UltiMate 3000 RSLCnano System attached to an Orbitrap Exploris 480 (Thermo Fisher Scientific). Peptides were injected on an Acclaim Pepmap trap column (Thermo Fisher Scientific #164564-CMD) before analysis on a PepMap RSLC C18 analytical column (Thermo Fisher Scientific #ES903) and eluted using a 125 min stepped gradient from 3 to 37% Buffer B (Buffer A: 0.1% formic acid in water, Buffer B: 0.08% formic acid in 80:20 acetonitrile:water (v:v)). Eluted peptides analyzed by the mass spectrometer operating in data independent acquisition (DIA) mode.

### MS data analysis

Peptide were searched against a human database containing isoforms (Uniprot Swissprot version release 05/10/2021) using DiaNN (v1.8.0) (Demichev et al., 2020) using library free mode. Statistical analysis was done in Perseus (v1.16.15.0) (Tyanova et al., 2016). Identified proteins with less than 2 unique peptides were excluded. Imputation of missing values was done using a gaussian distribution centered on the median with a downshift of 1.8 and width of 0.3, relative to the standard deviation, and intensities of proteins were median normalized. Significant changes between quadruplicate pulldowns of each chain type were assessed using ANOVA and P-values were adjusted using Benjamini Hochberg multiple hypothesis correction using a corrected P-value cutoff of <0.05. The list of 130 chain type specific binders was clustered using Spatial Hierarchical Euclidean clustering using the SciPy python library scipy.spatial.distance.pdist function (Virtanen et al., 2020) and visualized using the Plotly python library (Plotly Technologies Inc., 2015).

### Gene Ontology enrichment analysis

The DAVID web server (Sherman et al., 2022) was used for functional annotation and enrichment analyses. Enrichment of significant hits from the ANOVA of DIA MS pulldown with ubiquitin chains was analyzed against a background of all identified proteins. Annotation clusters linked to the six chain pulldown clusters were visualized using the Plotly python graphing library (Plotly Technologies Inc., 2015) and colored by DAVID enrichment score.

### UV-laser striping

U2OS cells stably expressing GFP-tagged fusions of NbSL3.3Q, NbSL18, DDB2 or GFP only under control of a Tetracycline promoter were seeded at approximately 10^5^ cells in 3.5 cm glass bottom dishes containing DMEM without phenol red supplemented with 10% FBS, 10 μM bromodeoxyuridine (BrdU, Sigma) and 1 μg/ml tetracycline (Sigma). UV laser micro-irradiation assays were performed at 37 °C and 5% CO2 atmosphere on a Leica TCS SP8X microscope system using a 355 nm coherent laser. Irradiation was applied in a bi-directional stripe pattern with an approximate 1.4-2.8J/m2 energy output. Images were acquired with a Leica HC PL APO CS2 63×/1.20 water objective operated by Leica LASX software in Live Data Mode. Images were acquired in autofocus mode starting with pre-irradiation state image and followed by a 10 min post-irradiation time-lapse. Images were stitched using an ImageJ macro and visualized using the Open Microscopy Environment Remote Objects (OMERO) server (Allan et al., 2012).

### Bioinformatics

Protein sequence alignments were generated using the EBI MAFFT server (https://www.ebi.ac.uk/Tools/msa/mafft/, Madeira *et al*., 2019) and secondary structures mapped with ENDscript 2 (http://endscript.ibcp.fr, Robert and Gouet, 2014). Binding site probabilities were predicted using the ScanNet web-server (http://bioinfo3d.cs.tau.ac.il/Scan-Net/, Tubiana, Schneidman-Duhovny and Wolfson, 2022).

## Supplemental information

**Table S1.** Crystallographic data collection and refinement statistics.

|  | K48-K63-Ub3 | NbSL3:K48-K63-Ub3 | NbSL3.3Q:K48-K63-Ub3 |
| --- | --- | --- | --- |
| **Data Collection** |  |  |  |
| Beamline | ID23-2, ESRF | ID23-2, ESRF | ID23-2, ESRF |
| Wavelength (Å) | 0.8731 | 0.8731 | 0.8731 |
| Space group | C2 | P1 21 1 | P1 |
| Total reflections | 19628 (1910) | 153451 (2434) | 176160 (12743) |
| Unique reflections | 9868 (976) | 79154 (1380) | 56422 (100) |
| a, b, c (Å) | 59.63 77.57 50.61 | 54.389 97.615 74.967 | 57.081 58.243 61.662 |
| $\alpha, \beta, \gamma$ (°) | 90.000 123.975 90.000 | 90 109.959 90 | 78.879 67.944 80.161 |
| Resolution (Å) | 24.73-2.19 (2.27-2.19) | 35.30-1.55 (1.61-1.55) | 40.86-1.86 (1.93-1.86) |
| Rmerge | 0.03991 (0.3488) | 0.06807 (0.5728) | 0.18 (1.447) |
| I/$\sigma$(I) | 8.80 (2.20) | 8.03 (1.60) | 4.96 (0.74) |
| Completeness (%) | 99.8 (98.68) | 74.33 (13.04) | 60.75 (1.66) |
| Multiplicity | 2.0 (2.0) | 1.9 (1.8) | 3.1 (3.1) |
| CC1/2 | 0.997 (0.843) | 0.992 (0.409) | 0.988 (0.285) |
| **Refinement** |  |  |  |
| R-work | 0.2239 (0.3573) | 0.1742 (0.2956) | 0.1866 (0.2481) |
| R-free | 0.2839 (0.4254) | 0.2123 (0.3477) | 0.2453 (0.2836) |
| **No. of Atoms** |  |  |  |
| Protein | 1774 | 5384 | 5445 |
| Ligand | 6 | 28 | 25 |
| Water | 25 | 722 | 523 |
| Wilson B-factor | 45.92 | 18.58 | 13.75 |
| **RMSDs** |  |  |  |
| Bond length (Å) | 0.003 | 0.007 | 0.002 |
| Bond angles (°) | 0.57 | 0.92 | 0.51 |
| **Ramachandran Plot (%)** |  |  |  |
| Favored region | 97.71 | 99.4 | 99.12 |
| Allowed region | 2.29 | 0.6 | 0.59 |
| Outlier region | 0 | 0 | 0.29 |
| PDB ID | 7NPO | 7NBB | 8A67 |

**Supplementary Figure 1.**
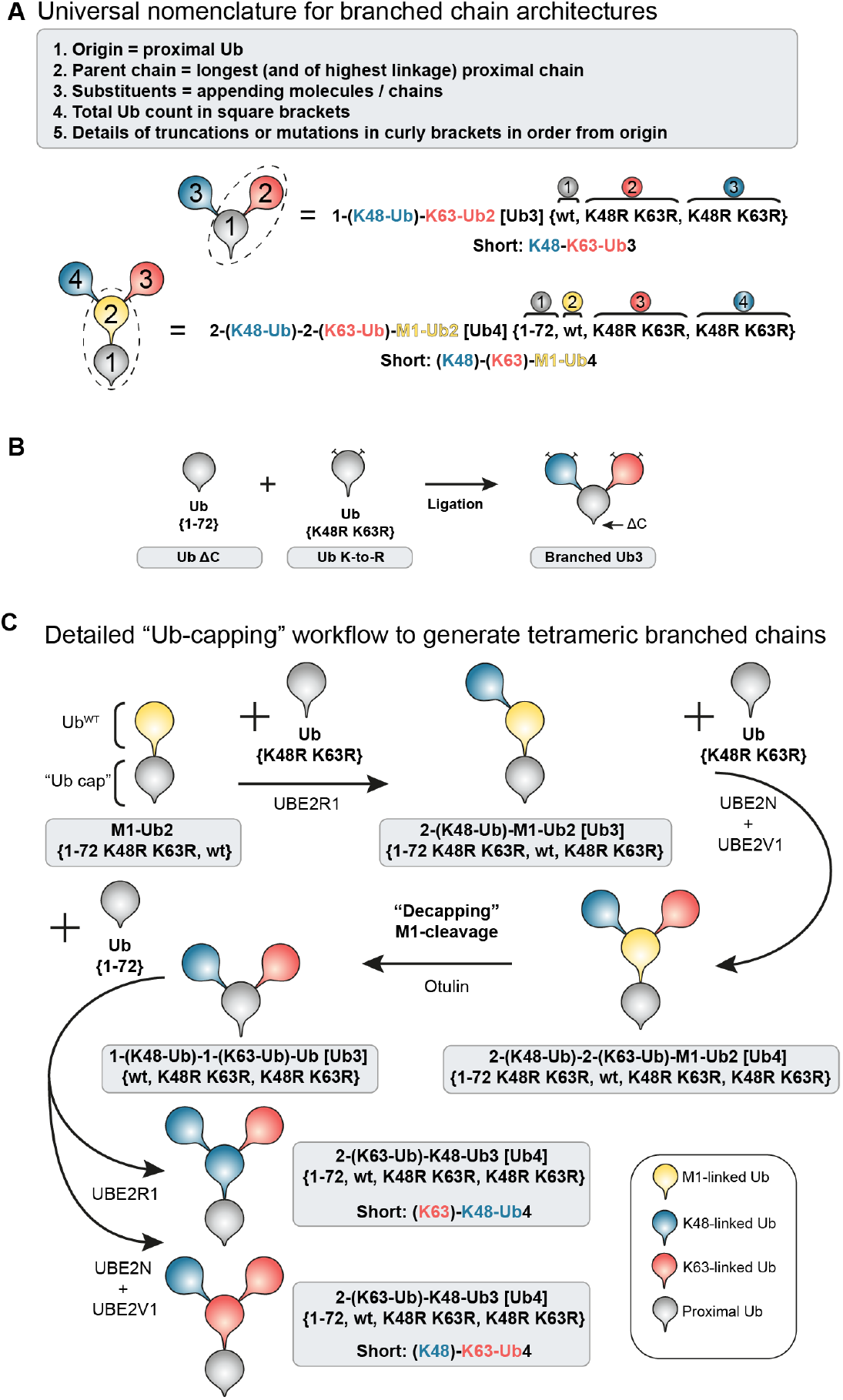
Nomenclature and detailed ligation of branched Ub chains. A) Rules of proposed Ub chain nomenclature to describe complex Ub chains and examples with chain schematics. Dotted circles indicate parent chain. B) ‘Delta-C’ ligation approach for generation of branched K48-K63-Ub3 chains. C) Detailed ‘Ub-capping’ workflow as shown in Figure 1B including full chain descriptions and ligation enzymes.

**Supplementary Figure 2.**
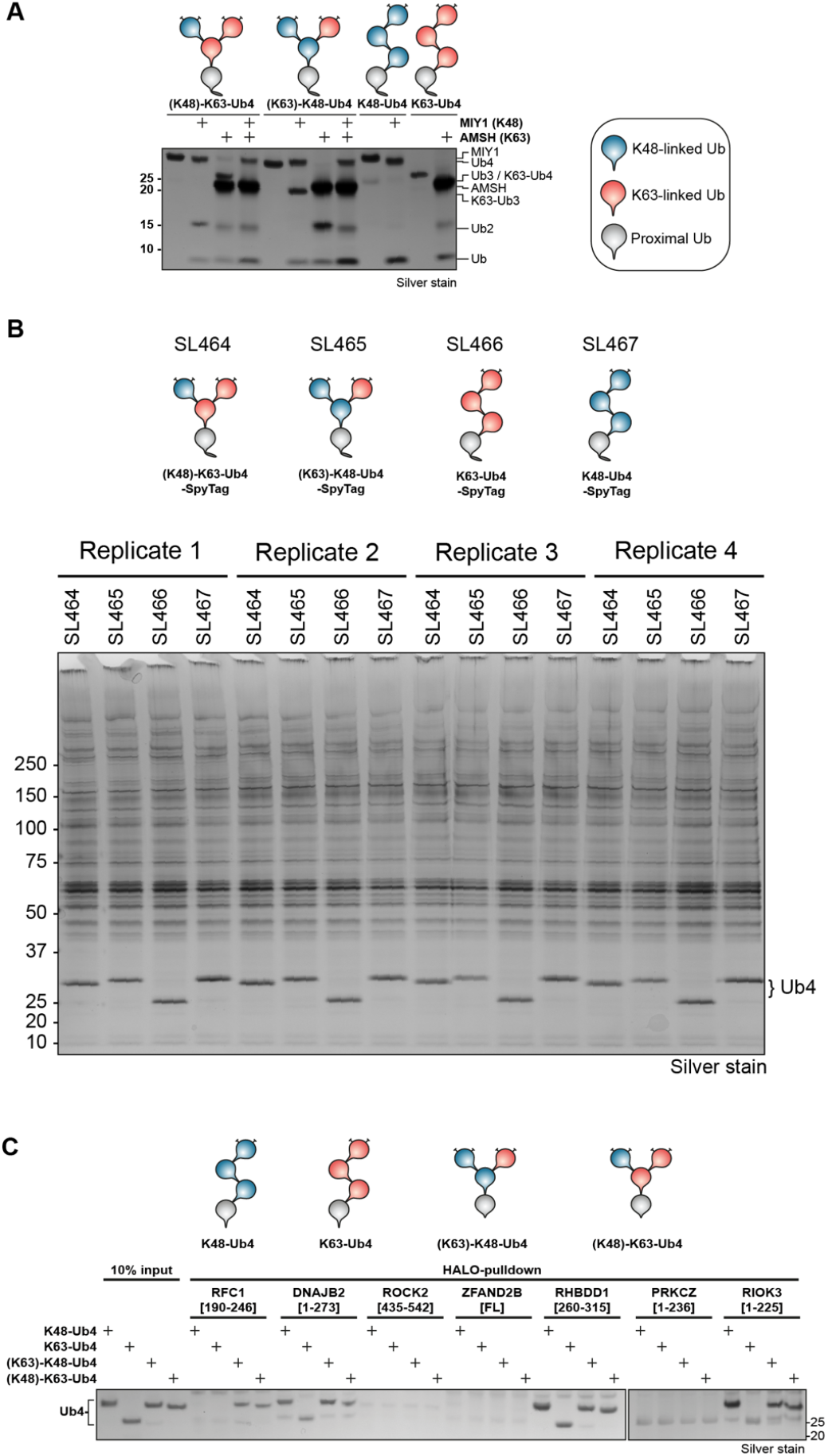
Quality control of chain pulldown and detailed HALO-pulldown with UBDs. A) Quality control DUB assay with linkage-specific enzymes Miy2/Ypl191c (K48) and AMSH (K63) of Spy-tagged branched and unbranched Ub4 chains. B) Silver-stained SDS-PAGE analysis of Ub chain pulldown samples. C) Silver-stained SDS-PAGE analysis of HALO pulldown with recombinant HALO-tagged UBDs and branched/un-branched Ub4 containing K48- and K63-linkages.

**Supplementary Figure 3.**
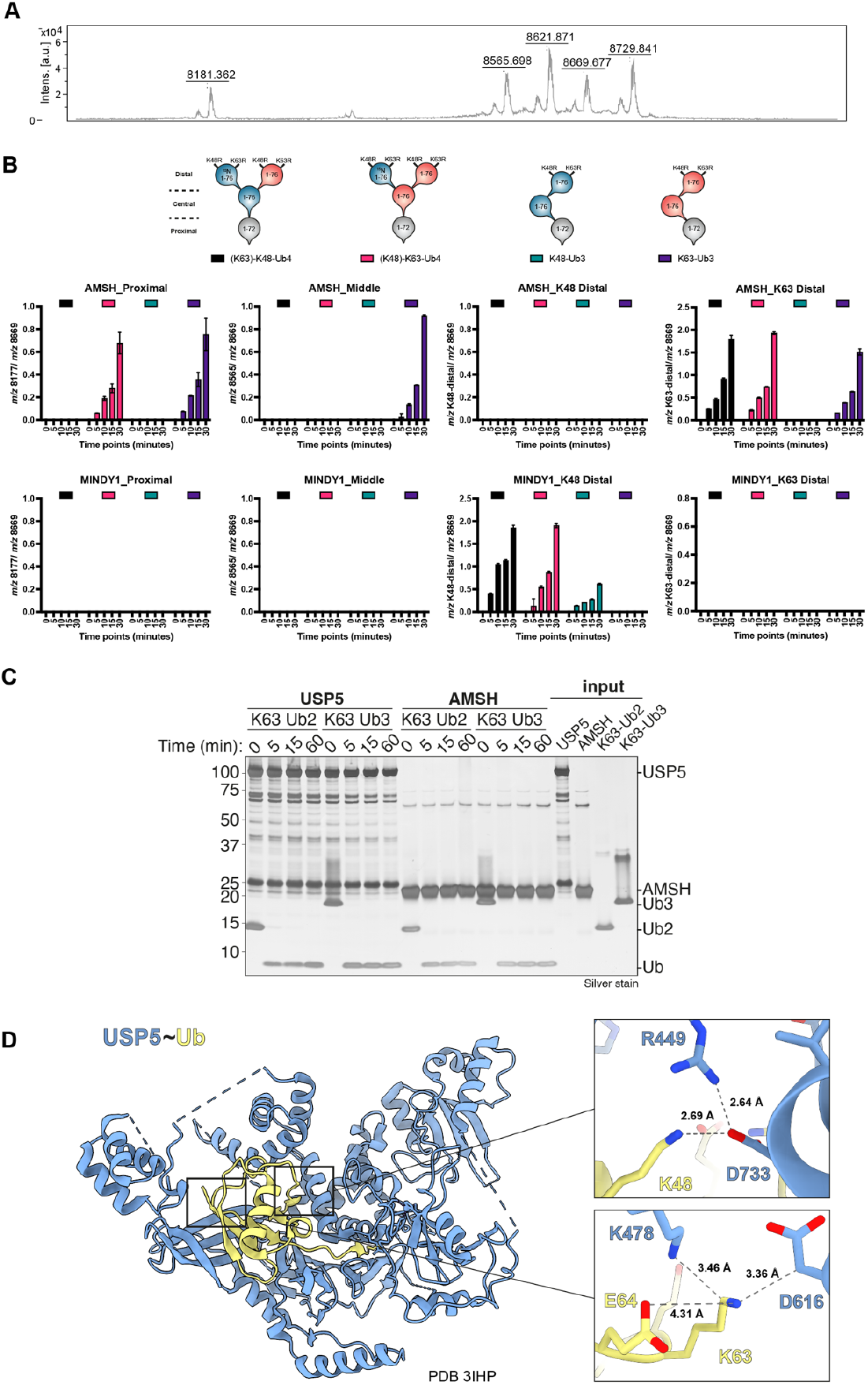
Establishing ULTIMAT DUB assay. A) Mass spectrum of fully cleaved Ub chain with indicated masses of the four released Ub moieties and added ^15^N-Ub internal standard. B) Initial quality control of ULTIMAT DUB assay with linkage-specific DUBs AMSH (K63) and MINDY1 (K48). Integrated mass peaks of released Ub moieties were normalized by ^15^N-Ub internal standard. C) Silver-stained SDS-PAGE analysis of DUB assay with USP5 and K63-specific DUB AMSH against K63-Ub2 and K63-Ub3 chains assembled from wild-type Ub. D) Crystal structure of catalytic domain of USP5 (blue) in complex with Ub (yellow) in cartoon representation (PDB 3IHP). Zoomed-in views of K48 and K63 residues of Ub and interacting USP5 residues as stick models with atomic distances indicated by dotted lines.

**Supplementary Figure 4.**
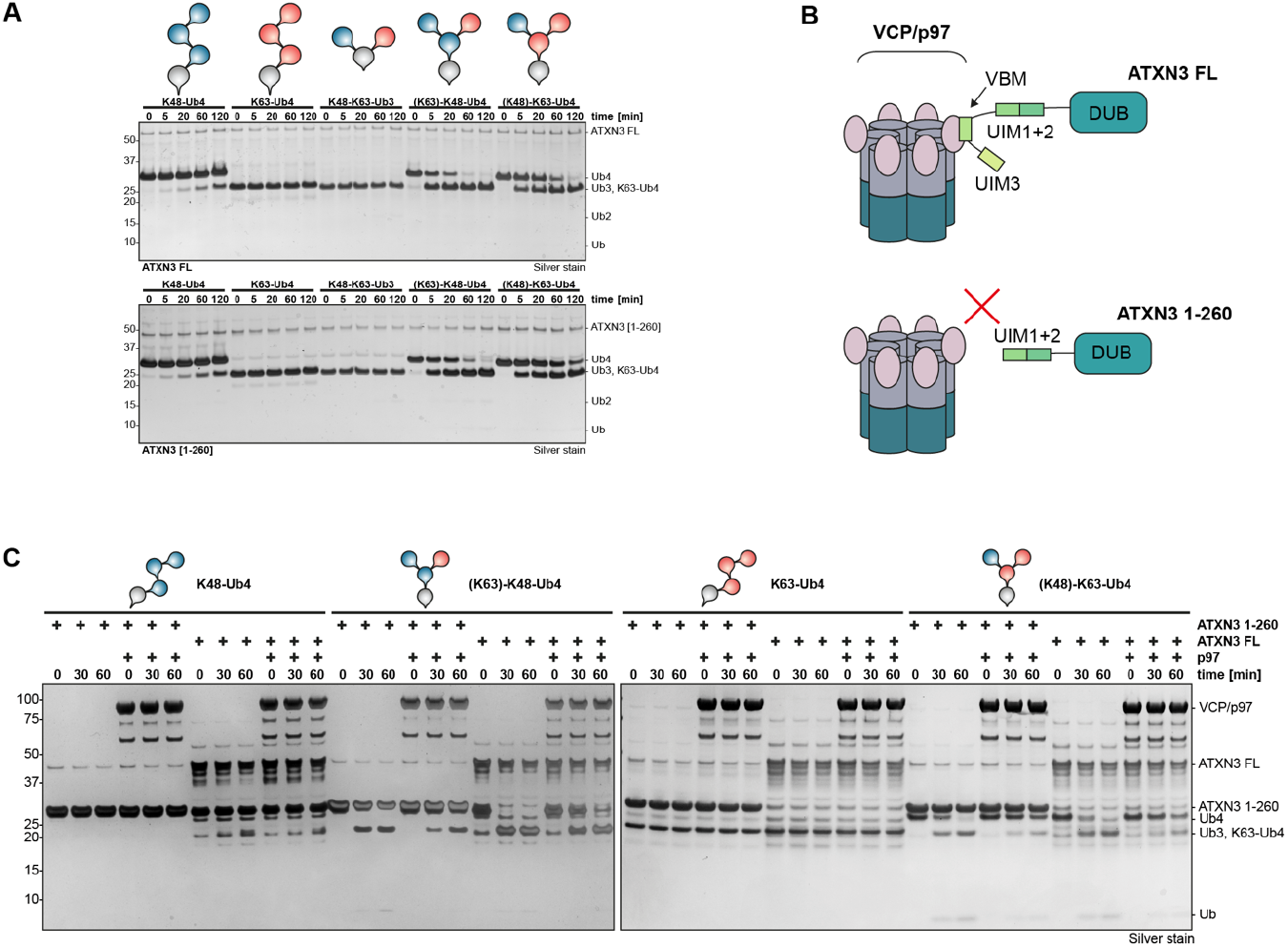
ATXN3 debranching activity is unaffected by recombinant p97 *in vitro*. A) Full gel of silver-stained SDS-PAGE analyses of DUB assays shown in Figure 4H. B) Schematic of VBM-mediated binding of full-length ATXN3 to VCP/p97 and truncated ATXN3 [1-260] lacking VMB. C) Silver-stained SDS-PAGE analyses of DUB assays of full-length ATXN3 and truncated ATXN3 [1-260] in the presence and absence of recombinant p97 against panel of branched and unbranched K48- and K63-linked Ub chains.

**Supplementary Figure 5.**
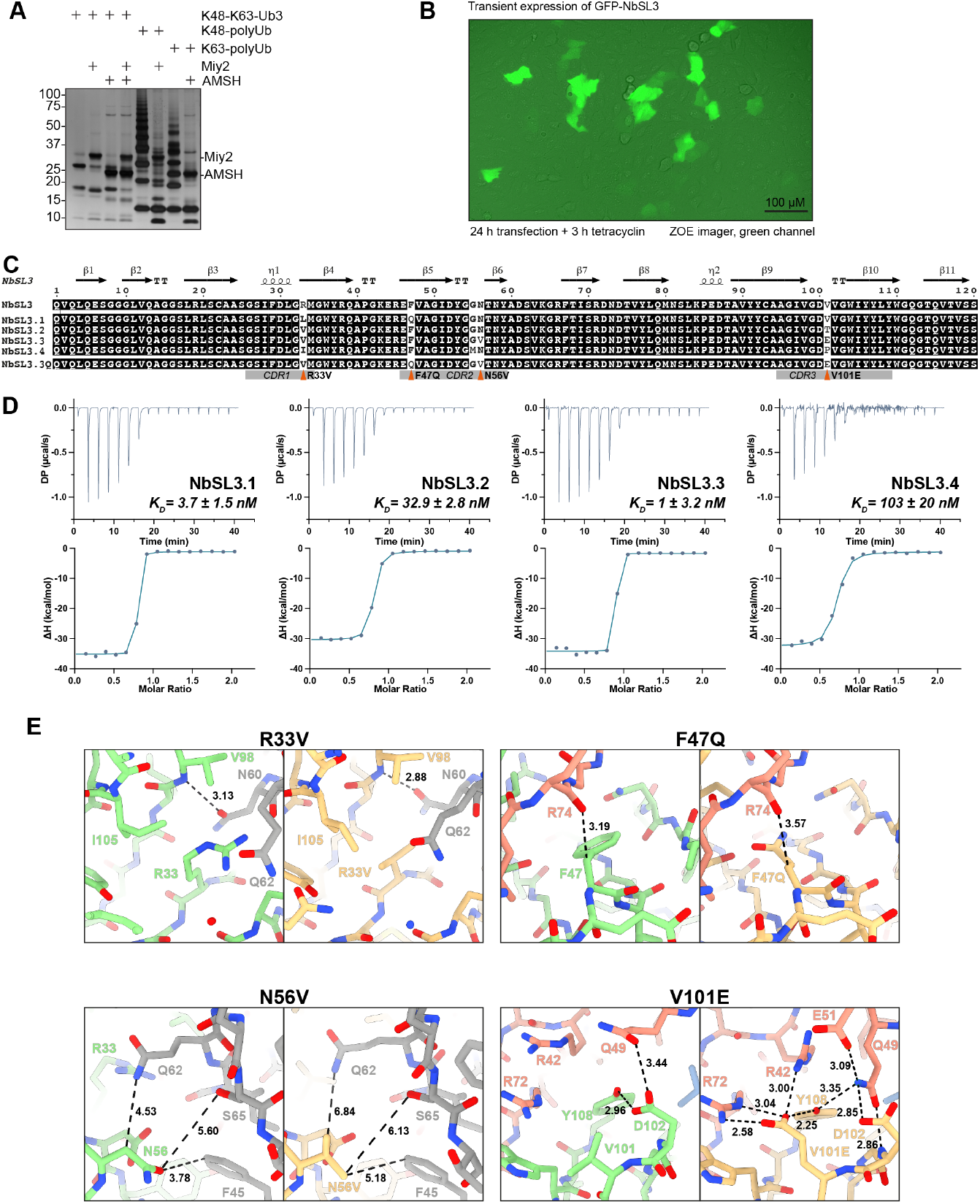
Selection and maturation of NbSL3.3Q. A) Silver-stained SDS-PAGE of DUB assay with AVI-tagged Ub chains used for nanobody selection and linkage-specific DUBs Miy2/Ypl191c (K48) and AMSH (K63). B) Live cell imaging of U2OS Flp-In cells transiently expressing NbSL3-GFP recorded using green channel of ZOE fluorescent cell imager. C) Sequence alignment, CDRs and secondary structure elements of NbSL3 and matured variants. The four mutations of the maturation from NbSL3 to NbSL3.3Q are indicated by red triangles. D) ITC analysis of second-generation NbSL3.1-4 binding to branched K48-K63-Ub3. E) Comparison of the residues affected by maturation mutations in the crystal structures of NbSL3 (green) and NbSL3.3Q (yellow) each in complex with branched K48-K63-Ub3 (K48-linked Ub in blue, K63-linked Ub in red, proximal Ub in grey). Distance measurements in Å indicated by black dotted lines.

**Supplementary Figure 6.**
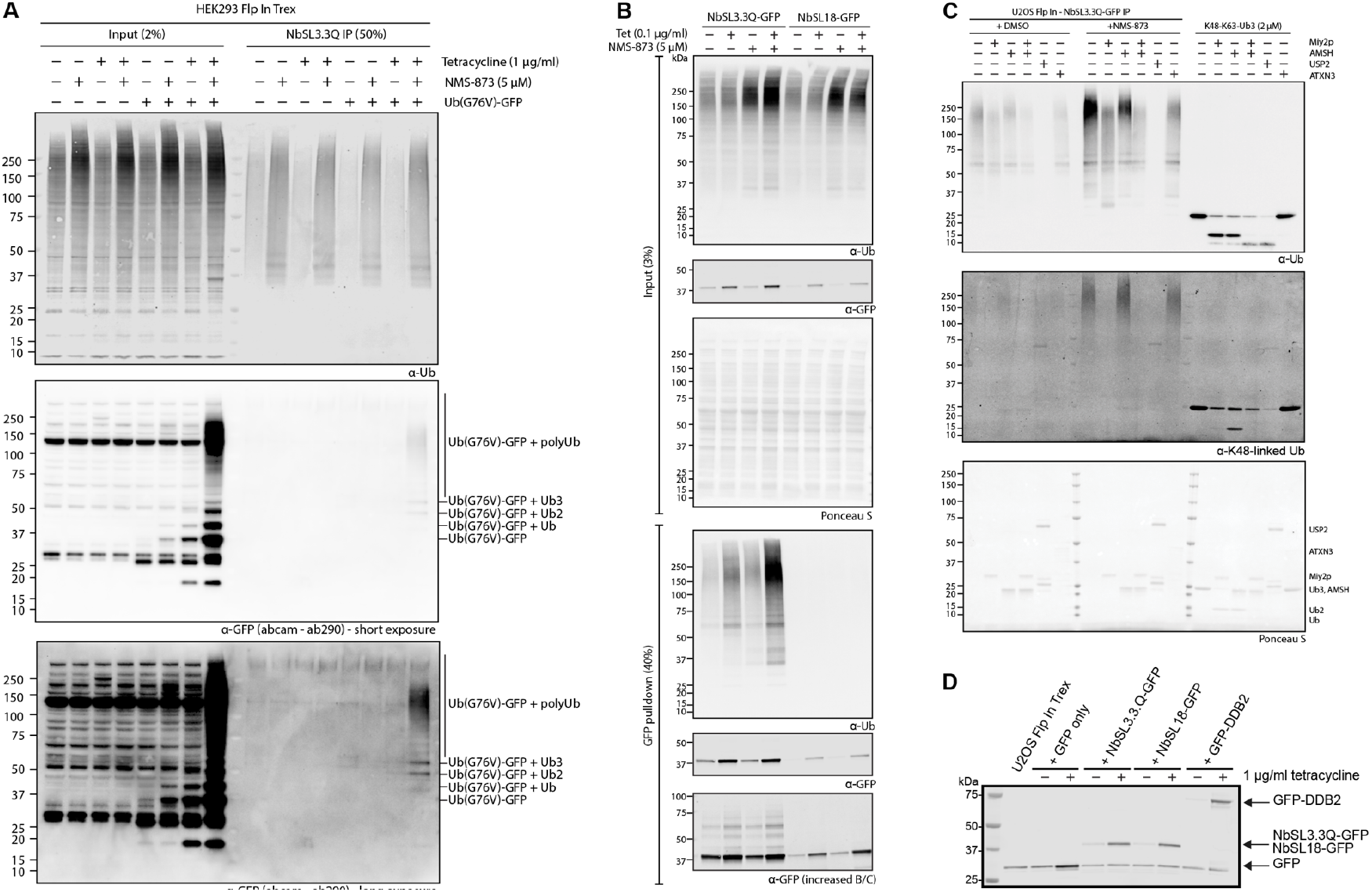
VCP/p97-substrate Ub^G76V^-GFP is modified with branched K48-K63-Ub and inhibition of VCP/p97 induces branching of K63-linked Ub off K48-chains. A) HEK293 Flp-In Trex cells were treated with tetracycline to induce expression of Ub^G76V^-GFP followed by VCP/p97-inhibition using NMS-873 (5 μM) for 4 hours. Subsequent pulldowns with NbSL3.3Q-immoblised agarose were analyzed by Western blotting for total Ub and GFP. Note: The anti-GFP antibody used here produces a non-specific band at approximately the same size as GFP. B) Anti-GFP pulldown from U2OS cells expressing either NbSL3.3Q-GFP or NbSL18-GFP following VCP/p97-inhibition with NMS-873 analyzed by Western blotting for total Ub and GFP. Note: The anti-GFP antibody used here produces a non-specific band at approximately the same size as GFP. C) DUB assay using linkage-specific DUBs Miy2/Ypl191c (K48) or AMSH (K63), non-specific DUB USP2 and debranching DUB ATXN3 (K63) incubated for 1 hour at 37°C with ubiquitin chains captured by anti-GFP pulldown from NbSL3.3Q-GFP expressing U2OS Flp-In Trex cells (lanes 1-12) following DMSO treatment (lanes 1-6) or VCP/p97-inhibition with NMS-873 (lanes 7-12), or recombinant branched K48-K63-Ub3 chains (lanes 13-18). Samples analyzed by Western blotting for total Ub and K48-linked Ub. D) Western blot analysis of anti-GFP pulldowns from U2OS Flp-In Trex used in UV micro-irradiation assay (Figure 6E) following tetracycline-induced expression of GFP, NbSL3.3Q-GFP, NbSL18-GFP or GFP-DDB2 visualized with anti-GFP antibody.

## References

Abdul Rehman, S.A., Armstrong, L.A., Lange, S.M., Kristariyanto, Y.A., Gräwert, T.W., Knebel, A., Svergun, D.I., Kulathu, Y., 2021. Mechanism of activation and regulation of deubiquitinase activity in MINDY1 and MINDY2. Molecular Cell 81, 4176–4190.e6.

Abdul Rehman, S.A., Kristariyanto, Y.A., Choi, S.-Y., Nkosi, P.J., Weidlich, S., Labib, K., Hofmann, K., Kulathu, Y., 2016. MINDY-1 Is a Member of an Evolutionarily Conserved and Structurally Distinct New Family of Deubiquitinating Enzymes. Molecular Cell 63, 146–155.

Ahlstedt, B.A., Ganji, R., Raman, M., 2022. The functional importance of VCP to maintaining cellular protein homeostasis. Biochemical Society Transactions BST20220648.

Akizuki, Y., Morita, M., Mori, Y., Kaiho-Soma, A., Dixit, S., Endo, A., Shimogawa, M., Hayashi, G., Naito, M., Okamoto, A., Tanaka, K., Saeki, Y., Ohtake, F., 2022. cIAP1-based degraders induce degradation via branched ubiquitin architectures. Nat Chem Biol 1–12.

Alexandru, G., Graumann, J., Smith, G.T., Kolawa, N.J., Fang, R., Deshaies, R.J., 2008. UBXD7 Binds Multiple Ubiquitin Ligases and Implicates p97 in HIF1α Turnover. Cell 134, 804–816.

Allan, C., Burel, J.-M., Moore, J., Blackburn, C., Linkert, M., Loynton, S., Macdonald, D., Moore, W.J., Neves, C., Patterson, A., Porter, M., Tarkowska, A., Loranger, B., Avondo, J., Lagerstedt, I., Lianas, L., Leo, S., Hands, K., Hay, R.T., Patwardhan, A., Best, C., Kleywegt, G.J., Zanetti, G., Swedlow, J. R., 2012. OMERO: flexible, model-driven data management for experimental biology. Nat Methods 9, 245–253.

Armstrong, L.A., Lange, S.M., Cesare, V.D., Matthews, S.P., Nirujogi, R.S., Cole, I., Hope, A., Cunningham, F., Toth, R., Mukherjee, R., Bojkova, D., Gruber, F., Gray, D., Wyatt, P.G., Cinatl, J., Dikic, I., Davies, P., Kulathu, Y., 2021. Biochemical characterization of protease activity of Nsp3 from SARS-CoV-2 and its inhibition by nanobodies. PLOS ONE 16, e0253364.

Azim, M., Surani, H., 1979. Glycoprotein synthesis and inhibition of glycosylation by tunicamycin in preimplantation mouse embryos: Compaction and trophoblast adhesion. Cell 18, 217–227.

Baranes-Bachar, K., Levy-Barda, A., Oehler, J., Reid, D.A., Soria-Bretones, I., Voss, T.C., Chung, D., Park, Y., Liu, C., Yoon, J.-B., Li, W., Dellaire, G., Misteli, T., Huertas, P., Rothenberg, E., Ramadan, K., Ziv, Y., Shiloh, Y., 2018. The Ubiquitin E3/E4 Ligase UBE4A Adjusts Protein Ubiquitylation and Accumulation at Sites of DNA Damage, Facilitating Double-Strand Break Repair. Molecular Cell 69, 866–878.e7.

Beck, D.B., Basar, M.A., Asmar, A.J., Thompson, J.J., Oda, H., Uehara, D.T., Saida, K., Pajusalu, S., Talvik, I., D’Souza, P., Bodurtha, J., Mu, W., Barañano, K.W., Miyake, N., Wang, R., Kempers, M., Tamada, T., Nishimura, Y., Okada, S., Kosho, T., Dale, R., Mitra, A., Macnamara, E., Undiagnosed Diseases Network, Matsumoto, N., Inazawa, J., Walkiewicz, M., Õunap, K., Tifft, C.J., Aksentijevich, I., Kastner, D.L., Rocha, P.P., Werner, A., 2021. Linkage-specific deubiquitylation by OTUD5 defines an embryonic pathway intolerant to genomic variation. Sci Adv 7, eabe2116.

Benatuil, L., Perez, J.M., Belk, J., Hsieh, C.-M., 2010. An improved yeast transformation method for the generation of very large human antibody libraries. Protein Engineering, Design and Selection 23, 155–159.

Beskow, A., Grimberg, K.B., Bott, L.C., Salomons, F.A., Dantuma, N.P., Young, P., 2009. A conserved unfoldase activity for the p97 AAA-ATPase in proteasomal degradation. J Mol Biol 394, 732–746.

Bett, J.S., Ritorto, M.S., Ewan, R., Jaffray, E.G., Virdee, S., Chin, J.W., Knebel, A., Kurz, T., Trost, M., Tatham, M.H., Hay, R.T., 2015. Ubiquitin C-terminal hydrolases cleave isopeptide-and peptide-linked ubiquitin from structured proteins but do not edit ubiquitin homopolymers. Biochem J 466, 489–498.

Boughton, A.J., Krueger, S., Fushman, D., 2020. Branching via K11 and K48 Bestows Ubiquitin Chains with a Unique Interdomain Interface and Enhanced Affinity for Proteasomal Subunit Rpn1. Structure 28, 29–43.e6.

Buchberger, A., Schindelin, H., Hänzelmann, P., 2015. Control of p97 function by cofactor binding. FEBS Letters, Dynamics, flexibility, and intrinsic disorder in protein assemblies 589, 2578–2589.

Chapple, J.P., Cheetham, M.E., 2003. The Chaperone Environment at the Cytoplasmic Face of the Endoplasmic Reticulum Can Modulate Rhodopsin Processing and Inclusion Formation *. Journal of Biological Chemistry 278, 19087–19094.

Cook, W.J., Jeffrey, L.C., Carson, M., Chen, Z., Pickart, C.M., 1992. Structure of a diubiquitin conjugate and a model for interaction with ubiquitin conjugating enzyme (E2). Journal of Biological Chemistry 267, 16467–16471.

Cooper, E.M., Cutcliffe, C., Kristiansen, T.Z., Pandey, A., Pickart, C.M., Cohen, R.E., 2009. K63-specific deubiquitination by two JAMM/MPN+ complexes: BRISC-associated Brcc36 and proteasomal Poh1. EMBO J 28, 621–631.

Custer, S.K., Neumann, M., Lu, H., Wright, A.C., Taylor, J.P., 2010. Transgenic mice expressing mutant forms VCP/p97 recapitulate the full spectrum of IBMPFD including degeneration in muscle, brain and bone. Hum Mol Genet 19, 1741–1755.

Davis, E.J., Lachaud, C., Appleton, P., Macartney, T.J., Näthke, I., Rouse, J., 2012. DVC1 (C1orf124) recruits the p97 protein segregase to sites of DNA damage. Nat Struct Mol Biol 19, 1093–1100.

De Cesare, V., Lopez, D.C., Mabbitt, P.D., Fletcher, A.J., Soetens, M., Antico, O., Wood, N.T., Virdee, S., 2021. Deubiquitinating enzyme amino acid profiling reveals a class of ubiquitin esterases. Proceedings of the National Academy of Sciences 118.

De Cesare, V., Moran, J., Traynor, R., Knebel, A., Ritorto, M.S., Trost, M., McLauchlan, H., Hastie, C.J., Davies, P., 2020. High-throughput matrix-assisted laser desorption/ionization time-of-flight (MALDI-TOF) mass spectrometry-based deubiquitylating enzyme assay for drug discovery. Nat Protoc 15, 4034–4057.

Demichev, V., Messner, C.B., Vernardis, S.I., Lilley, K.S., Ralser, M., 2020. DIA-NN: neural networks and interference correction enable deep proteome coverage in high throughput. Nat Methods 17, 41–44.

Deol, K.K., Crowe, S.O., Du, J., Bisbee, H.A., Guenette, R.G., Strieter, E.R., 2020. Proteasome-Bound UCH37/UCHL5 Debranches Ubiquitin Chains to Promote Degradation. Molecular Cell.

Du, J., Babik, S., Li, Y., Deol, K.K., Eyles, S.J., Fejzo, J., Tonelli, M., Strieter, E., 2022. A cryptic K48 ubiquitin chain binding site on UCH37 is required for its role in proteasomal degradation. eLife 11, e76100.

El Oualid, F., Merkx, R., Ekkebus, R., Hameed, D.S., Smit, J.J., de Jong, A., Hilkmann, H., Sixma, T.K., Ovaa, H., 2010. Chemical Synthesis of Ubiquitin, Ubiquitin-Based Probes, and Diubiquitin. Angewandte Chemie International Edition 49, 10149–10153.

Emmerich, C.H., Ordureau, A., Strickson, S., Arthur, J.S.C., Pedrioli, P.G.A., Komander, D., Cohen, P., 2013. Activation of the canonical IKK complex by K63/M1-linked hybrid ubiquitin chains. Proceedings of the National Academy of Sciences of the United States of America 110, 15247–52.

Erpapazoglou, Z., Dhaoui, M., Pantazopoulou, M., Giordano, F., Mari, M., Léon, S., Raposo, G., Reggiori, F., Haguenauer-Tsapis, R., 2012. A dual role for K63-linked ubiquitin chains in multivesicular body biogenesis and cargo sorting. Mol Biol Cell 23, 2170–2183.

Faesen, A.C., Luna-Vargas, M.P.A., Geurink, P.P., Clerici, M., Merkx, R., van Dijk, W.J., Hameed, D.S., El Oualid, F., Ovaa, H., Sixma, T.K., 2011. The differential modulation of USP activity by internal regulatory domains, interactors and eight ubiquitin chain types. Chem Biol 18, 1550–1561.

Fernández-Sáiz, V., Buchberger, A., 2010. Imbalances in p97 co-factor interactions in human proteinopathy. EMBO Rep 11, 479–485.

Fleig, L., Bergbold, N., Sahasrabudhe, P., Geiger, B., Kaltak, L., Lemberg, M.K., 2012. Ubiquitin-Dependent Intramembrane Rhomboid Protease Promotes ERAD of Membrane Proteins. Molecular Cell 47, 558–569.

Fottner, M., Brunner, A.-D., Bittl, V., Horn-Ghetko, D., Jussupow, A., Kaila, V.R.I., Bremm, A., Lang, K., 2019. Site-specific ubiquitylation and SUMOylation using genetic-code expansion and sortase. Nat Chem Biol 15, 276–284.

French, M.E., Koehler, C.F., Hunter, T., 2021. Emerging functions of branched ubiquitin chains. Cell Discov 7, 1–10.

Fujisawa, R., Polo Rivera, C., Labib, K.P., 2022. Multiple UBX proteins reduce the ubiquitin threshold of the mammalian p97-UFD1-NPL4 unfoldase. eLife 11, e76763.

Gaidt, M.M., Morrow, A., Fairgrieve, M.R., Karr, J.P., Yosef, N., Vance, R.E., 2021. Self-guarding of MORC3 enables virulence factor-triggered immunity. Nature 600, 138–142.

Gitlin, A.D., Heger, K., Schubert, A.F., Reja, R., Yan, D., Pham, V.C., Suto, E., Zhang, J., Kwon, Y.C., Freund, E.C., Kang, J., Pham, A., Caothien, R., Bacarro, N., Hinkle, T., Xu, M., McKenzie, B.S., Haley, B., Lee, W.P., Lill, J.R., Roose-Girma, M., Dohse, M., Webster, J.D., Newton, K., Dixit, V.M., 2020. Integration of innate immune signalling by caspase-8 cleavage of N4BP1. Nature 587, 275–280.

Haakonsen, D.L., Rape, M., 2019. Branching Out: Improved Signaling by Heterotypic Ubiquitin Chains., Trends in cell biology. Elsevier.

Hermanns, T., Pichlo, C., Woiwode, I., Klopffleisch, K., Witting, K.F., Ovaa, H., Baumann, U., Hofmann, K., 2018. A family of unconventional deubiquitinases with modular chain specificity determinants. Nature Communications 9, 799.

Hou, P., Yang, K., Jia, P., Liu, L., Lin, Y., Li, Z., Li, J., Chen, S., Guo, S., Pan, J., Wu, J., Peng, H., Zeng, W., Li, C., Liu, Y., Guo, D., 2021. A novel selective autophagy receptor, CCDC50, delivers K63 polyubiquitination-activated RIG-I/MDA5 for degradation during viral infection. Cell Res 31, 62–79.

Husnjak, K., Dikic, I., 2012. Ubiquitin-Binding Proteins: Decoders of Ubiquitin-Mediated Cellular Functions. Annual Review of Biochemistry 81, 291–322.

Ito, A., Takahashi, R., Miura, C., Baba, Y., 1975. Synthetic Study of Peptide Aldehydes. Chemical & Pharmaceutical Bulletin 23, 3106–3113.

Johnson, E.S., Ma, P.C.M., Ota, I.M., Varshavsky, A., 1995. A Proteolytic Pathway That Recognizes Ubiquitin as a Degradation Signal *. Journal of Biological Chemistry 270, 17442–17456.

Keeble, A.H., Banerjee, A., Ferla, M.P., Reddington, S.C., Anuar, I.N.A.K., Howarth, M., 2017. Evolving Accelerated Amidation by SpyTag/SpyCatcher to Analyze Membrane Dynamics. Angewandte Chemie - International Edition 56, 16521–16525.

Keusekotten, K., Elliott, P.R., Glockner, L., Fiil, B.K., Damgaard, R.B., Kulathu, Y., Wauer, T., Hospenthal, M.K., Gyrd-Hansen, M., Krappmann, D., Hofmann, K., Komander, D., 2013. OTULIN Antagonizes LUBAC Signaling by Specifically Hydrolyzing Met1-Linked Polyubiquitin. Cell 153, 1312–1326.

Komander, D., Rape, M., 2012. The Ubiquitin Code. Annual Review of Biochemistry 81, 203–229.

Kondo, H., Rabouille, C., Newman, R., Levine, T.P., Pappin, D., Freemont, P., Warren, G., 1997. p47 is a cofactor for p97-mediated membrane fusion. Nature 388, 75–78.

Krissinel, E., 2015. Stock-based detection of protein oligomeric states in jsPISA. Nucleic Acids Research 43, W314–W319.

Kristariyanto, Y.A., Abdul Rehman, S.A., Weidlich, S., Knebel, A., Kulathu, Y., 2017. A single MIU motif of MINDY-1 recognizes K48-linked polyubiquitin chains. EMBO reports 18, 392–402.

Kwasna, D., Abdul Rehman, S.A., Natarajan, J., Matthews, S., Madden, R., De Cesare, V., Weidlich, S., Virdee, S., Ahel, I., Gibbs-Seymour, I., Kulathu, Y., 2018. Discovery and Characterization of ZUFSP/ZUP1, a Distinct Deubiquitinase Class Important for Genome Stability. Molecular Cell 70, 150–164.e6.

Laço, M.N., Cortes, L., Travis, S.M., Paulson, H.L., Rego, A.C., 2012. Valosin-containing protein (VCP/p97) is an activator of wild-type ataxin-3. PLoS One 7, e43563.

Lange, S.M., Armstrong, L.A., Kulathu, Y., 2022. Deubiquitinases: From mechanisms to their inhibition by small molecules. Molecular Cell 82, 15–29.

Lauwers, E., Jacob, C., André, B., 2009. K63-linked ubiquitin chains as a specific signal for protein sorting into the multivesicular body pathway. Journal of Cell Biology 185, 493–502.

Lee, D.H., Goldberg, A.L., 1998. Proteasome inhibitors: valuable new tools for cell biologists. Trends in Cell Biology 8, 397–403.

Li, S., Paulsson, K.M., Chen, S., Sjögren, H.-O., Wang, P., 2000. Tapasin Is Required for Efficient Peptide Binding to Transporter Associated with Antigen Processing *. Journal of Biological Chemistry 275, 1581–1586.

Long, L., Thelen, J.P., Furgason, M., Haj-Yahya, M., Brik, A., Cheng, D., Peng, J., Yao, T., 2014. The U4/U6 recycling factor SART3 has histone chaperone activity and associates with USP15 to regulate H2B deubiquitination. J Biol Chem 289, 8916–8930.

Madeira, F., Park, Y.M., Lee, J., Buso, N., Gur, T., Madhusoodanan, N., Basutkar, P., Tivey, A.R.N., Potter, S.C., Finn, R.D., Lopez, R., 2019. The EMBL-EBI search and sequence analysis tools APIs in 2019. Nucleic acids research.

Magnaghi, P., D’Alessio, R., Valsasina, B., Avanzi, N., Rizzi, S., Asa, D., Gasparri, F., Cozzi, L., Cucchi, U., Orrenius, C., Polucci, P., Ballinari, D., Per-rera, C., Leone, A., Cervi, G., Casale, E., Xiao, Y., Wong, C., Anderson, D.J., Galvani, A., Donati, D., O’Brien, T., Jackson, P.K., Isacchi, A., 2013. Covalent and allosteric inhibitors of the ATPase VCP/p97 induce cancer cell death. Nat Chem Biol 9, 548–556.

Manglik, A., Kobilka, B.K., Steyaert, J., 2017. Nanobodies to Study G Protein-Coupled Receptor Structure and Function. Annu Rev Pharmacol Toxicol 57, 19–37.

McMahon, C., Baier, A.S., Pascolutti, R., Wegrecki, M., Zheng, S., Ong, J.X., Erlandson, S.C., Hilger, D., Rasmussen, S.G.F., Ring, A.M., Manglik, A., Kruse, A.C., 2018. Yeast surface display platform for rapid discovery of conformationally selective nanobodies. Nature Structural and Molecular Biology 25, 289–296.

McNeill, H., Knebel, A., Arthur, J.S.C., Cuenda, A., Cohen, P., 2004. A novel UBA and UBX domain protein that binds polyubiquitin and VCP and is a substrate for SAPKs. Biochemical Journal 384, 391–400.

Meerang, M., Ritz, D., Paliwal, S., Garajova, Z., Bosshard, M., Mailand, N., Janscak, P., Hübscher, U., Meyer, H., Ramadan, K., 2011. The ubiquitin-selective segregase VCP/p97 orchestrates the response to DNA double-strand breaks. Nat Cell Biol 13, 1376–1382.

Messick, T.E., Greenberg, R.A., 2009. The ubiquitin landscape at DNA double-strand breaks. Journal of Cell Biology 187, 319–326.

Mevissen, T.E.T., Hospenthal, M.K., Geurink, P.P., Elliott, P.R., Akutsu, M., Arnaudo, N., Ekkebus, R., Kulathu, Y., Wauer, T., El Oualid, F., Freund, S.M.V., Ovaa, H., Komander, D., 2013. OTU Deubiquitinases Reveal Mechanisms of Linkage Specificity and Enable Ubiquitin Chain Restriction Analysis. Cell 154, 169–184.

Meyer, H., Weihl, C.C., 2014. The VCP/p97 system at a glance: connecting cellular function to disease pathogenesis. J Cell Sci 127, 3877–3883.

Meyer, H.H., Shorter, J.G., Seemann, J., Pappin, D., Warren, G., 2000. A complex of mammalian Ufd1 and Npl4 links the AAA-ATPase, p97, to ubiquitin and nuclear transport pathways. The EMBO Journal 19, 2181–2192.

Meyer, H.-J., Rape, M., 2014. Enhanced protein degradation by branched ubiquitin chains. Cell 157, 910–921.

Nakasone, M.A., Livnat-Levanon, N., Glickman, M.H., Cohen, R.E., Fushman, D., 2013. Mixed-Linkage Ubiquitin Chains Send Mixed Messages. Structure 21, 727–740.

Nathan, J.A., Tae Kim, H., Ting, L., Gygi, S.P., Goldberg, A.L., 2013. Why do cellular proteins linked to K63-polyubiquitin chains not associate with proteasomes? EMBO J 32, 552–565.

Nepravishta, R., Ferrentino, F., Mandaliti, W., Mattioni, A., Weber, J., Polo, S., Castagnoli, L., Cesareni, G., Paci, M., Santonico, E., 2019. CoCUN, a Novel Ubiquitin Binding Domain Identified in N4BP1. Biomolecules 9, 284.

Ohtake, F., 2022. Branched ubiquitin code: from basic biology to targeted protein degradation. The Journal of Biochemistry 171, 361–366.

Ohtake, F., Saeki, Y., Ishido, S., Kanno, J., Tanaka, K., 2016. The K48-K63 Branched Ubiquitin Chain Regulates NF-κB Signaling. Molecular Cell 64, 251–266.

Ohtake, F., Tsuchiya, H., Saeki, Y., Tanaka, K., 2018. K63 ubiquitylation triggers proteasomal degradation by seeding branched ubiquitin chains. Proceedings of the National Academy of Sciences of the United States of America 115, E1401–E1408.

Panier, S., Durocher, D., 2009. Regulatory ubiquitylation in response to DNA double-strand breaks. DNA Repair, Ubiquitin, SUMO and the Maintenance of Genome Stability 8, 436–443.

Patterson-Fortin, J., Shao, G., Bretscher, H., Messick, T.E., Greenberg, R.A., 2010. Differential regulation of JAMM domain deubiquitinating enzyme activity within the RAP80 complex. J Biol Chem 285, 30971–30981.

Pawson, T., 2004. Specificity in signal transduction: from phosphotyrosine-SH2 domain interactions to complex cellular systems. Cell 116, 191–203.

Plotly Technologies Inc., 2015. Collaborative data science [WWW Document]. URL https://plot.ly

Podust, L.M., Podust, V.N., Sogo, J.M., Hübscher, U., 1995. Mammalian DNA polymerase auxiliary proteins: analysis of replication factor C-catalyzed proliferating cell nuclear antigen loading onto circular double-stranded DNA. Molecular and Cellular Biology 15, 3072–3081.

Pruneda, J.N., Komander, D., 2019. Chapter Fourteen - Evaluating enzyme activities and structures of DUBs, in: Hochstrasser, M. (Ed.), Methods in Enzymology, Ubiquitin and Ubiquitin-like Protein Modifiers. Academic Press, pp. 321–341.

Raasi, S., Orlov, I., Fleming, K.G., Pickart, C.M., 2004. Binding of polyubiquitin chains to ubiquitin-associated (UBA) domains of HHR23A. J Mol Biol 341, 1367–1379.

Raman, M., Sergeev, M., Garnaas, M., Lydeard, J.R., Huttlin, E.L., Goessling, W., Shah, J.V., Harper, J.W., 2015. Systematic proteomics of the VCP-UBXD adaptor network identifies a role for UBXN10 in regulating ciliogenesis. Nat Cell Biol 17, 1356–1369.

Rao, M.V., Williams, D.R., Cocklin, S., Loll, P.J., 2017. Interaction between the AAA+ ATPase p97 and its cofactor ataxin3 in health and disease: Nu-cleotide-induced conformational changes regulate cofactor binding. Journal of Biological Chemistry 292, 18392–18407.

Ritorto, M.S., Ewan, R., Perez-Oliva, A.B., Knebel, A., Buhrlage, S.J., Wightman, M., Kelly, S.M., Wood, N.T., Virdee, S., Gray, N.S., Morrice, N.A., Alessi, D.R., Trost, M., 2014. Screening of DUB activity and specificity by MALDI-TOF mass spectrometry. Nature Communications 5, 4763.

Robert, X., Gouet, P., 2014. Deciphering key features in protein structures with the new ENDscript server. Nucleic Acids Research 42, W320–W324.

Sherman, B.T., Hao, M., Qiu, J., Jiao, X., Baseler, M.W., Lane, H.C., Imamichi, T., Chang, W., 2022. DAVID: a web server for functional enrichment analysis and functional annotation of gene lists (2021 update). Nucleic Acids Res gkac194.

Sims, J.J., Cohen, R.E., 2009. Linkage-specific avidity defines the lysine 63-linked polyubiquitin-binding preference of rap80. Mol Cell 33, 775–783.

Singh, A.N., Oehler, J., Torrecilla, I., Kilgas, S., Li, S., Vaz, B., Guérillon, C., Fielden, J., Hernandez-Carralero, E., Cabrera, E., Tullis, I.D., Meerang, M., Barber, P.R., Freire, R., Parsons, J., Vojnovic, B., Kiltie, A.E., Mailand, N., Ramadan, K., 2019. The p97-Ataxin 3 complex regulates homeostasis of the DNA damage response E3 ubiquitin ligase RNF8. The EMBO Journal 38, e102361.

Sobhian, B., Shao, G., Lilli, D.R., Culhane, A.C., Moreau, L.A., Xia, B., Livingston, D.M., Greenberg, R.A., 2007. RAP80 Targets BRCA1 to Specific Ubiquitin Structures at DNA Damage Sites. Science 316, 1198–1202.

Song, A., Hazlett, Z., Abeykoon, D., Dortch, J., Dillon, A., Curtiss, J., Martinez, S.B., Hill, C.P., Yu, C., Huang, L., Fushman, D., Cohen, R.E., Yao, T., 2021. Branched ubiquitin chain binding and deubiquitination by UCH37 facilitate proteasome clearance of stress-induced inclusions. eLife 10, e72798.

Swatek, K.N., Usher, J.L., Kueck, A.F., Gladkova, C., Mevissen, T.E.T., Pruneda, J.N., Skern, T., Komander, D., 2019. Insights into ubiquitin chain architecture using Ub-clipping. Nature 572, 533–537.

Tubiana, J., Schneidman-Duhovny, D., Wolfson, H.J., 2022. ScanNet: an interpretable geometric deep learning model for structure-based protein binding site prediction. Nat Methods 19, 730–739.

Tyanova, S., Temu, T., Sinitcyn, P., Carlson, A., Hein, M.Y., Geiger, T., Mann, M., Cox, J., 2016. The Perseus computational platform for comprehensive analysis of (prote)omics data. Nat Methods 13, 731–740.

van den Boom, J., Meyer, H., 2017. VCP/p97-Mediated Unfolding as a Principle in Protein Homeostasis and Signaling. Molecular Cell 69, 182–194.

Virtanen, P., Gommers, R., Oliphant, T.E., Haberland, M., Reddy, T., Cournapeau, D., Burovski, E., Peterson, P., Weckesser, W., Bright, J., van der Walt, S.J., Brett, M., Wilson, J., Millman, K.J., Mayorov, N., Nelson, A.R.J., Jones, E., Kern, R., Larson, E., Carey, C.J., Polat, I., Feng, Y., Moore, E.W., VanderPlas, J., Laxalde, D., Perktold, J., Cimrman, R., Henriksen, I., Quintero, E.A., Harris, C.R., Archibald, A.M., Ribeiro, A.H., Pedregosa, F., van Mulbregt, P., 2020. SciPy 1.0: fundamental algorithms for scientific computing in Python. Nat Methods 17, 261–272.

Wang, C., Deng, L., Hong, M., Akkaraju, G.R., Inoue, J., Chen, Z.J., 2001. TAK1 is a ubiquitin-dependent kinase of MKK and IKK. Nature 412, 346–351.

Weeks, S.D., Grasty, K.C., Hernandez-Cuebas, L., Loll, P.J., 2009. Crystal structures of Lys-63-linked tri-and di-ubiquitin reveal a highly extended chain architecture. Proteins: Structure, Function, and Bioinformatics 77, 753–759.

Williamson, D.S., Borgognoni, J., Clay, A., Daniels, Z., Dokurno, P., Drysdale, M. J., Foloppe, N., Francis, G.L., Graham, C.J., Howes, R., Macias, A.T., Murray, J.B., Parsons, R., Shaw, T., Surgenor, A.E., Terry, L., Wang, Y., Wood, M., Massey, A.J., 2009. Novel Adenosine-Derived Inhibitors of 70 kDa Heat Shock Protein, Discovered Through Structure-Based Design. J. Med. Chem. 52, 1510–1513.

Winborn, B.J., Travis, S.M., Todi, S.V., Scaglione, K.M., Xu, P., Williams, A.J., Cohen, R.E., Peng, J., Paulson, H.L., 2008. The Deubiquitinating Enzyme Ataxin-3, a Polyglutamine Disease Protein, Edits Lys63 Linkages in Mixed Linkage Ubiquitin Chains*. Journal of Biological Chemistry 283, 26436–26443.

Yau, R.G., Doerner, K., Castellanos, E.R., Haakonsen, D.L., Werner, A., Wang, N., Yang, X.W., Martinez-Martin, N., Matsumoto, M.L., Dixit, V.M., Rape, M., 2017. Assembly and Function of Heterotypic Ubiquitin Chains in Cell-Cycle and Protein Quality Control. Cell 171, 918–933.e20.

Zhang, X., Smits, A.H., van Tilburg, G.B.A., Jansen, P.W.T.C., Makowski, M.M., Ovaa, H., Vermeulen, M., 2017. An Interaction Landscape of Ubiquitin Signaling. Mol Cell 65, 941–955.e8.

Zhang, Y., Klein, B.J., Cox, K.L., Bertulat, B., Tencer, A.H., Holden, M.R., Wright, G.M., Black, J., Cardoso, M.C., Poirier, M.G., Kutateladze, T.G., 2019. Mechanism for autoinhibition and activation of the MORC3 ATPase. Proc Natl Acad Sci U S A 116, 6111–6119.

Zhong, X., Pittman, R.N., 2006. Ataxin-3 binds VCP/p97 and regulates retrotranslocation of ERAD substrates. Hum Mol Genet 15, 2409–2420.

